# Minimal epistatic networks from integrated sequence and mutational protein data

**DOI:** 10.1101/2023.09.25.559251

**Authors:** Simona Cocco, Lorenzo Posani, Rémi Monasson

## Abstract

Predicting the functional effects of mutations to a wild-type protein sequence is a major computational challenge. We introduce here a computationally efficient procedure to identify the few, most informative epistatic links between residues in a protein, integrating sequence data and functional measurements with mutational scans. Our approach shows performances comparable to state-of-the-art deep networks, while requiring much less parameters and being hence much more interpretable. The selected network links mostly focus on the protein functional sites, adapt to the *in vitro* or *in vivo* function experimentally tested, and are not necessary related to structural contacts.

## 1. Introduction

Much progress has been made in inferring protein structure from homologous sequence data over recent years [1, 2, 3, 4]. One of the simplest approaches is the graphical Potts model (also called Direct Coupling Analysis, DCA), in which residues directly interact through epistatic couplings inferred to reproduce the covariation between amino acids [5, 6, 7, 8, 9, 10, 11]. Large couplings have been shown to predict contacts between residues in the tridimensional structure of the protein, and could therefore be used to guide protein [12, 13, 14, 15, 16] and protein complexes [17, 18, 19] predictions.

Apart from protein folding, a challenge in evolutionary biology and bioinformatics is to predict the effect of mutations from a wild-type sequence. Progress along this direction would be important for medical applications, e.g. in predicting antibiotic resistance [20, 21, 22, 23], in designing drugs against rapidly evolving viruses, such as HIV [24, 25, 26] and SARS-Cov2 [27, 28]. Recent developments of deep scanning mutagenesis experiments combined with the advances in sequencing technology have nowadays shifted the paradigm from few tested mutations to large mutational scans, eg. by testing almost all single site mutations around a wild type sequence [22, 20, 21, 29, 30, 31, 32, 33, 34, 35, 36, 37, 38, 39, 40, 41, 42, 43, 44], single and double mutations in a small protein region[44], or mutational paths on some chosen sites and amino-acids [45].

A variety of models and bioinformatics tools have been used to score the effect of amino-acid substitutions. A first class of such models is trained on sequence data collected for a protein family after aligning them in a Multi Sequence Alignements (MSA), and has a large spectrum of approaches going from the simplest independent-site models based on position weight matrices [46, 47] eventually combined with phylogenetic informations [48, 49], to Graphical Potts models including epistatic interactions [24, 23, 50, 49, 48], deep neural networks [51]. A second class includes language model learned more generally on all available sequence data (Uniprot), eventually combined with mutational data [52, 53, 54, 55]. A third class includes models trained from structural data [56, 57, 58].

Sequence-based models have been shown to be able to predict the effect of amino-acid substitutions on a protein property, measured through mutational scans [59, 60, 61]. However, it remains unclear why and to what extent homologous sequences, which reflect the complex pattern of structural and functional constraints that have shaped organisms throughout evolutionary times can be informative about a specific biochemical property. This question is particularly difficult, due to the multiple aspects of functionalities performed by proteins (stability, affinity, specificity, catalytic activity, allostery, …), resulting in a variety of constraints on their sequences. The presence within the same protein family of subfamilies, each characterized by different functional specificities, is well known, and occurs at the level of the organism (phylogeny), or the gene location, e.g. for multidomains with slightly different roles [21]. Disentangling such complexity and focusing on the functionality that is experimentally investigated in a mutational scan is challenging, especially for families with evolutionary diverged proteins. Principal component analysis and clustering approaches have been used to separate subfamilies, with different functionalities and to identify key residues for the protein function [62, 63, 64, 65, 30, 66, 11, 32].

The aim of the present work is to introduce a procedure to infer functionality-specific minimal graphical models using sequence and partial single mutation scans. The models are minimal in the sense that as little epistatic links between residues as possible are introduced. This approach ensures robustness (due to the implicit regularization resulting from the high level of sparsity of the interaction graph) and facilitates the interpretability of the few present links. Our procedure allows us to precisely predict mutational effects for new data and to identify the core network for the tested functionality. The number of links is determined as a function of the quality and the quantity of available data [67].

The paper is organized as follows. We first define the *K*-link Potts model, aiming at predicting the effects of mutations on a protein sequence of length *N* from a minimal epistatic network between residues, containing only 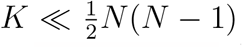 links. We then introduce MutInf, a mutational scan informed procedure, able to iteratively improve the choice of the *K* most relevant links. The performance of our approach is assessed on 10 mutational data sets [20, 21, 22, 29, 30, 33, 34, 39, 41, 42], and is cross-validated on test data sets, including separate single and double site mutations. We investigate in particular how performances depend on the initial ranking of couplings, and reach saturation as the MutInf updating procedure is iterated. We then analyze and discuss the optimal functional networks, identify the most connected sites, and study how the inferred functional networks change with the selective pressure applied in the experimental setting. Two illustrations are presented: *β*lactamase, for which mutational effects under varying ampicillin concentration have been experimentally investigated in [21], and the Ubiquitin protein, for which mutational effects have been measured for the overall fitness and for the binding of a particular ligand (E1) in [41] and [42]. Conclusions and perspectives are reported in the Discussion section.

### 2. Results

Our modeling approach, based on the inference of a sparse graphical model for a protein sequence (of length *N*), is schematized in Fig. 1. Briefly speaking, a ranking heuristics initially produces an ordered list of all putative epistatic links. The parameters of the model, couplings and local fields, are inferred from sequence data. The model is then used to predict the effects of point mutations on a wild-type sequence. Confrontation with experimental mutational scan data allows for updating the ranking of the most relevant

**Figure 1.**
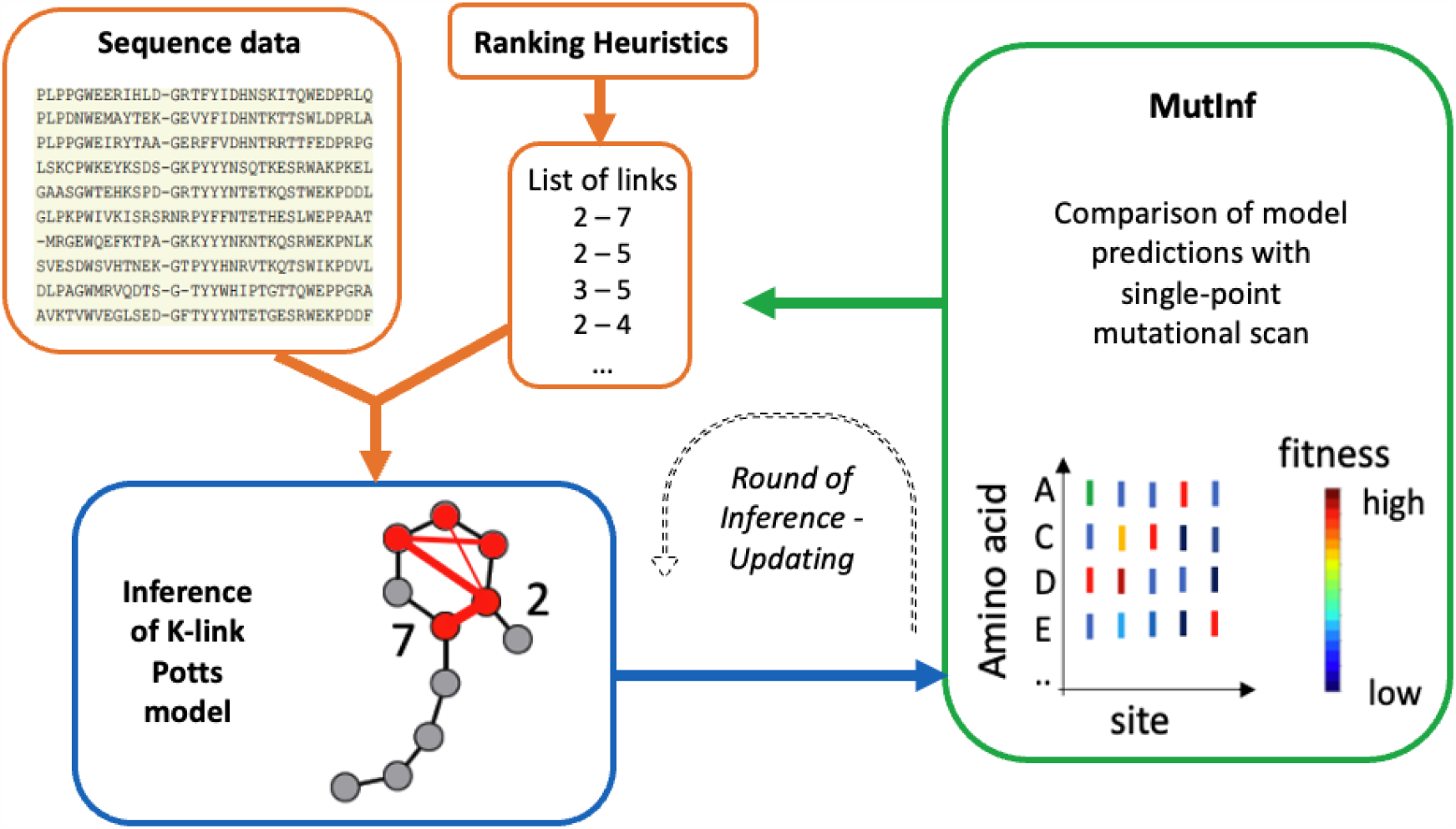
Pipeline for the inference of the *K*-link Potts model (blue box) based on multi-sequence alignment data and a heuristic ranking of the relevant epistatic links (orange boxes). The MutInf procedure updates the ranking of couplings through a comparison of the model predictions and mutation scan data (green box). The whole procedure can be iterated for several rounds, until convergence is reached. The number of couplings associated to the best performance determines *K*.

links and prune the other ones. The procedure can be iterated for several rounds, until it converges to an optimal sparse model. We now describe these steps in more detail.

### plmDCA-based initial list of links (p heuristic)

We first need as input an ordered list of all 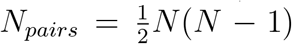 links. This ranking will decide in which order links are progressively included in our model. The first heuristic we have used, referred to as *p* hereafter, is based on plmDCA: A fully connected Potts model is first inferred with the standard Pseudo-Likelihood Method, also called plmDCA [10, 50], using the available sequence data (see Suppl. Appendix A.4 for details and choice of regularization). Pairs (*i, j*) are then ranked in decreasing order of the Frobenius norms of their coupling matrices. We will discuss alternative heuristics in the following.

### Inference of K-link Potts model

Given a list of *K* links we infer from sequence (MSA) data a *K*-Link Potts model. The parameters of the *K*-Link Potts model are: (1) the local fields (Position Weight Matrices) *h*_*i*_(*a*), where *a* is the amino acid on site *i*, (2) the *K* epistatic coupling matrices *J*_*k*_(*a, b*) for the selected links (*k* = 1 … *K*), where *a, b* are the amino acids carried by the linked residues, respectively, *i*(*k*), *j*(*k*). The values of the fields and of the *K* coupling parameters are computed through a fast and accurate approximation scheme, see Methods and [68, 69]. Once the parameters have been inferred the predictor of the fitness cost resulting from the mutation → *wt*_*i*_ *a, i*.*e*. the change of amino acid on the site *i* of the wild-type (*wt*) sequence to *a*, reads

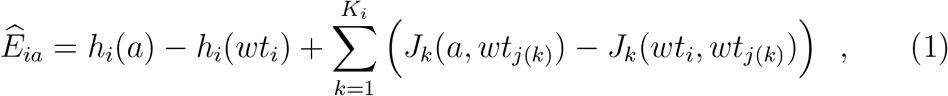

where the sums runs on the *K*_*i*_ links connecting the site *i* to the neighbour sites *j*. This neighborhood may be empty (*K*_*i*_ = 0) for some sites.

### Mutation-Informed (MutInf) updating procedure

We infer all the *K*-link Potts models, for *K* = 0 (independent sites) to *N*_*pairs*_ (fully connected network), adding on link at the time, following the initial ranking of the links. For each *K*, we compute the Spearman correlation *Sp*_*K*_ between the estimators 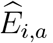 and the experimental measurements *E*_*ia*_ of the fitness changes across all single-point mutations. *Sp*_*K*_ is a measure of the predictive power of the corresponding *K*-links Potts model.

The difference 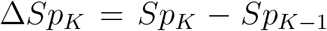 is an estimate of how much the *K*^*th*^ link, say, (*i, j*), contributes to the Spearman correlation. It can be positive (the addition of the link to the network of epistatic couplings improve predictive performance), or negative (performance are lowered by the insertion of the link); notice that the value of 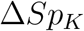 depends on the *K* −1 links preceding its insertion, and not only on the sites *i, j* it connects. Ordering all the links according to their contributions Δ*Sp* results in a new ranking of the pairs of sites. The procedure of inference of *K*-links Potts models then resumes according to this new ranking, see Fig. 1.

The two phases of link ranking and network parameter inference can be repeated in successive rounds, until the *Sp* reaches convergence. The value of *K* corresponding to the highest *Sp* value defines the degree of sparsity of the optimal *K*-link Potts model at each round. The MutInf procedure is sped up by pruning at each round links with negative Spearman contributions Δ*Sp*, making the list of possible links smaller and smaller rounds after rounds.

### Optimal performance and convergence of the MutInf procedure

We show in Fig. 2A the Spearman (*Sp*) and Pearson (*R*^2^) correlations between the costs of single mutations around the *wt* sequence of the RNA binding domain of Saccharomyces cerevisiae predicted by the *K*-link model and their experimental counterparts, estimated from the relative growth rates of the strains [39]. The initial rankings of links was done with the *p* heuristic; other choices are discussed below. At the initial iteration, denoted by round 0 *r* = 0 (green curve in Fig. 2A), we observe that both correlations strongly vary, and show a non-monotonic behaviour with *K*. Not surprisingly, the independent-site model (*K* = 0, olive circles in Fig. 2A) gives poorest performance. *Sp* and *R*^2^ then increase as epistatic links are progressively introduced, reaching a maximal value. For larger *K*, performance decreases due to overfitting of the sequence data; note that for very dense networks, our inference approximation scheme is not expected to be accurate any longer.The *K*-link model with maximal Spearman correlation (green stars in Fig. 2A) shows performances comparable to the ones of a fully-connected model inferred by plmDCA under strong regularization on the coupling parameters (square symbols, see Methods Sec.4.4) [70]. This is a remarkable result as the optima 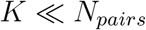, and the network is highly sparse.

**Figure 2.**
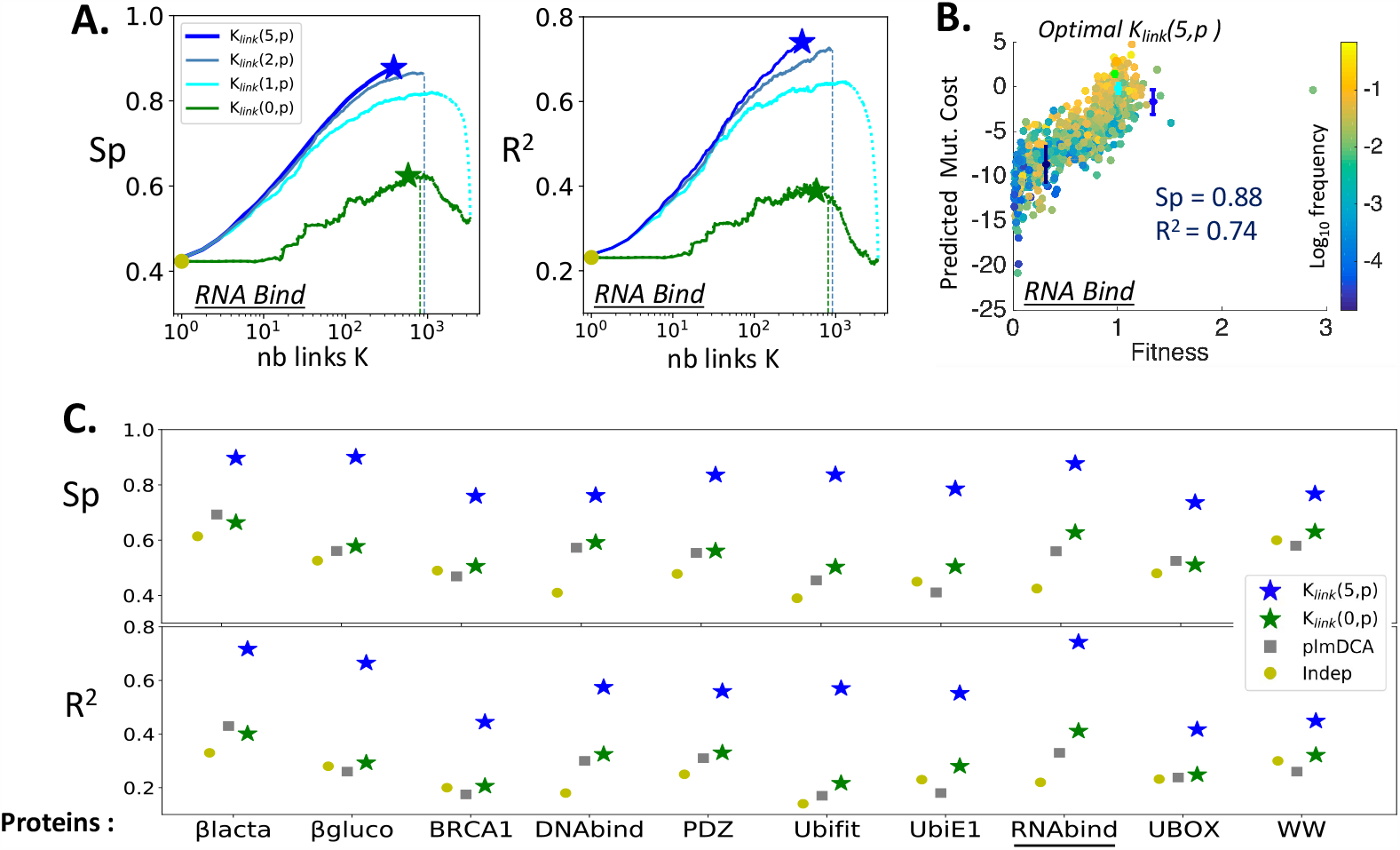
Performances of the *K*-link Potts model inferred with the MutInf procedure for predicting single mutation effects. *K*_*link*_(*r, p*) refers to the optimal *K*-link Potts model after *r* rounds of MutInf with heuristics *p* (plmDCA ranking). **A**. Spearman (*Sp*) and Pearson (*R*^2^) correlations between the predicted and measured fitness effects resulting from single mutations of the RNA binding domain (*N* = 82 amino acids) of the wild-type yeast strain [39] vs. number *K* of links in the Potts model. Green, cyan, gray, blue curves: rounds *r* = 0, 1, 2, 5 of MutInf. Only links with positive Spearman contributions are kept after each round. Stars indicate the optimal Spearman at rounds *r* = 0 and *r* = 5. Vertical lines: theoretical cut-off, from the signal to noise ratio criterion, for rounds 0 and 2 and for the maximal number of links in the graph. **B**. Scatter plots of predicted vs. experimental (log-enrichment ratios with respect to *wt* yeast strain [39]) costs of mutations for the optimal network (blue star in panel A). Each mutation is shown as a dot, colors represent the log-frequencies (base 10) of the mutations in the sequence data. Statistical error bars over *E*_*ia*_ are shown for 9 mutations, randomly chosen at different frequencies; 4 of them are smaller than the dot size. **C**. *Sp* and *R*^2^ for all 10 mutational scans under study (*N* varying between 31 and 441, see Methods, Table 1), according to an independent-site (Indep), plmDCA, and *K*-link Potts models, and for plmDCA initial ranking heuristic.

After one re-ranking by MutInf, the performance largely increases, compare curves corresponding to *r* = 1 (cyan curve) to *r* = 0 rounds in Fig. 2A. Due to the pruning of links having negative Spearman contribution at each round of MutInf (see Methods Sec. 4.6), the effective number of links rapidly diminishes. Optimal performance and network sparsity do not vary any longer after *r* = 5 rounds. The optimal network is very sparse (blue star in Fig. 2A), with *K* = 387 selected links corresponding to ≃ 11% of the *N*_*pairs*_ pairs of sites. Fig. 2B shows that the scatter plot of fitness predictions and measurements for the optimal network, with correlation coefficients *Sp* = 0.88 and *R*^2^ = 0.74.

**Table 1:** List of protein datasets. Numbers *N* of sites, *B* of sequences (after removing redundant sequences in the alignment), fraction *f* of measured single mutations over all possible ones (19*N*), and Spearman correlation 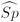 between experimental measurements and Potts model predictions from [50].

| Protein Name (wild-type) | $N$ | $B$ | $f$ | $\widehat{Sp}$ |
| --- | --- | --- | --- | --- |
| βlactamase (BLAT-ECOLX) | 253 | 7620 | 0.96 | 0.7 |
| β-Glucosidase (BG) | 441 | 26606 | 0.6 | 0.62 |
| DNA Binding domain (GAL4-YEAST) | 63 | 11278 | 1 | 0.6 |
| PDZ (DLG4-RAT) | 84 | 24795 | 0.99 | 0.5 |
| RNA Binding (PABP-YEAST) | 82 | 70779 | 0.76 | 0.6 |
| Ring Domain (BRCA1-HUMAN ) | 75 | 14448 | 0.97 | 0.47 |
| WW (YAP1-HUMAN) | 31 | 8251 | 0.54 | 0.57 |
| Ubiquitin(RL401_YEAST) ftn | 71 | 10023 | 0.86 | 0.42 |
| Ubiquitin(RL401_YEAST) E1 | 71 | 10023 | 0.96 | 0.42 |
| UBOX Domain (UBE48-MOUSE) | 76 | 6153 | 0.85 | 0.52 |

The above study was repeated for the 10 mutational scans over 9 proteins under study, with results reported in Fig. 2C. For all protein families we observe a substantial increase in performance after few rounds of MutInf reranking, with Spearman correlations increasing from ≃0.4 −0.6 for *r* = 0 to ≃0.8 − 0.9 for *r* = 5 rounds. As shown in Suppl. Fig. C.8 *Sp* converges in 1-5 rounds depending on the protein family and, more specifically, on its length and the improvement in *Sp* with *r* is accompanied by an increase of *R*^2^, *i*.*e*. by a closer to linear relationship between the predicted and the experimental fitness costs.

### Estimation of the network sparsity level depending on the number of data

A theoretical estimate of the maximal number of links that the inference procedure can account for without overfitting the data is shown in Fig. 2A (and Suppl. Fig. C.8 for all the cases under study); this estimate closely matches the number of links providing maximal performances.

Our estimate is based on theoretical expressions for the variances of the predicted mutational costs (Methods, Section 4.4). These variances depend on the number of sequence data in the MSA used to fit the model, on the site-dependent frequencies of the wild-type and mutated amino acids in the MSA, see Fig. 2B, and increase with the numbers *K*_*i*_ of links from sites *i* in Eq. 1, see Methods Eq. 9 and Suppl. Fig. C.9. We then estimate the maximal number of links in the network as the value of *K* at which the theoretical standard deviation for the single prediction become comparable to the overall range of variation of predicted mutational costs [67], see Methods Section 4.5 and Suppl. Fig. C.9. This estimate of the maximal *K*, averaged over the 10 mutational scans, leads to only 0.2% of Spearman loss with respect to the optimum found through a systematic scan of *K* values, see Fig. 2B and Suppl. Table B.2, Suppl. Fig. C.8.

### Predictive power of the K-link Potts model

We then assess the predictive power of the optimal *K*-link Potts model identified above on three different types of test sets, independent from the data used to train the model: *i)* different experimental settings for the mutational scans on the same wild type and a largely overlapping set of single site mutations, *ii)* single-site mutations not included in the training set, *iii)* some tested double mutations (recall the training set include single-site mutations only).

### Test on different single mutation scans on the same protein: the βlactamase case

In Fig. 3A we select the MutInf optimal network based on the mutational scan data of Firnberg et al. [22] and it on the experimental data by Stiffler et al. [21] ^1^. For both datasets the *wt* protein is the E. Coli *β*lactamase and the fitness effects of almost all single-site mutations were experimentally tested. The mutational scans are however different: Firnberg et al. measure the fitness through the bacterial growth rate, while Stiffler et al. estimate the fitness based on the *β*lactamase resistance in a cellular synthetic-biology device. Fig. 3A shows equivalent performances on the test and training data sets, with a Spearman coefficients *Sp* ≃ 0.88. The same performances are obtained when exchanging the training and test data sets.

**Figure 3.**
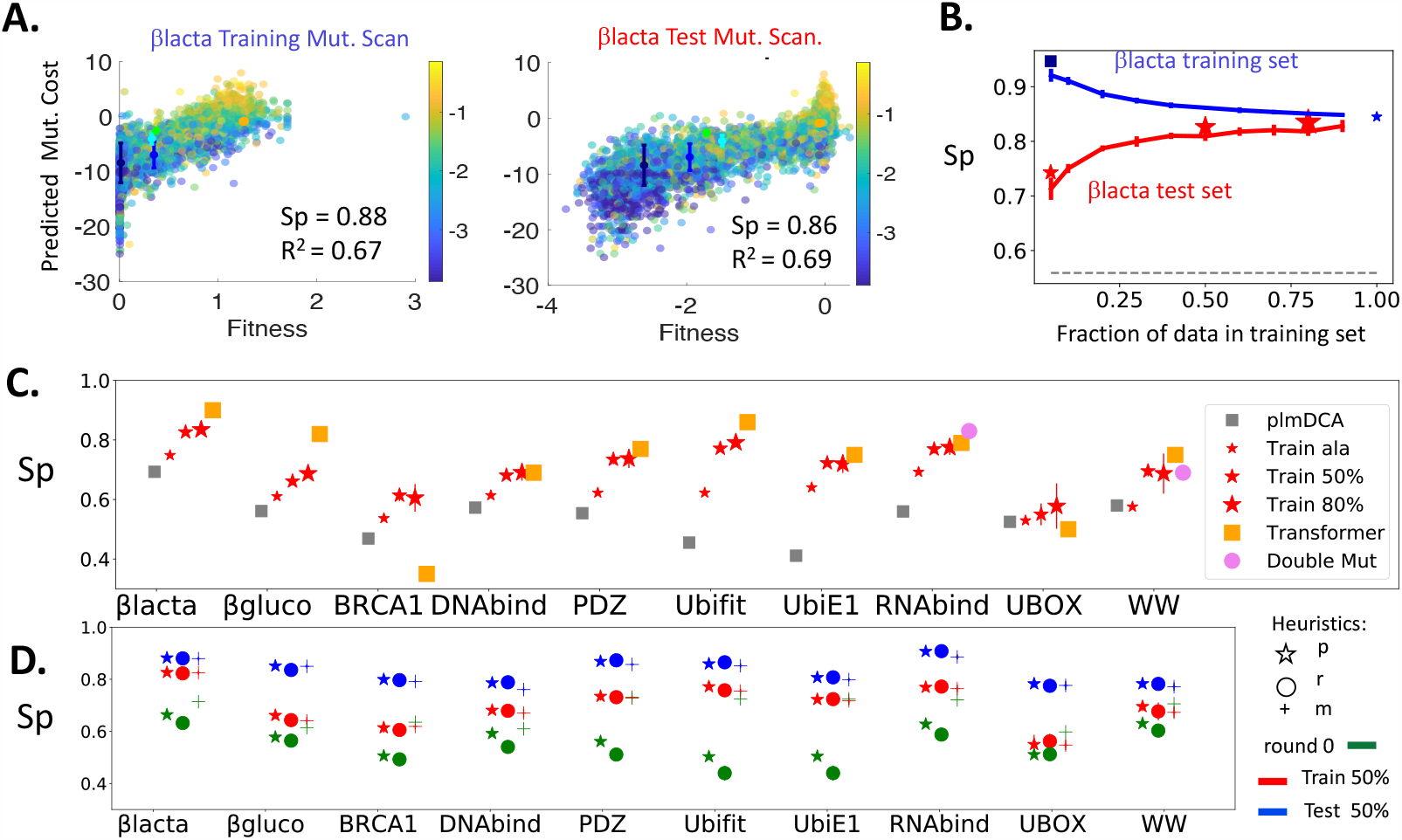
Cross-validation of the *K*-link model with the MutInf selection procedure. **A**. Different single mutational scan on *β*lactamase, same wild type (TEM1). Fitnesses (y-axis) predicted by *K*_*link*_(4, *p*) trained on Firnberg et al.’s dataset [22] vs. results of the single point mutational scans of Firnberg et al. (left) and Stiffler et al. [21] (right); Spearman between the two data sets: 0.94 (Suppl. Fig.C.11). Similar results obtained by exchanging the train and test data sets are shown in Suppl. Fig.C.10. **B**. Cross-validation of *K*_*link*_(2, *p*) trained on random partitions of the data by Firnberg et al. Results show the Spearman *Sp* on the training and test sets as functions of the relative size of the former; bars indicate standard deviations over 10 repetitions. A similar behavior is observed for *R*^2^, see Suppl. Fig. C.13. **C**. Cross-validation over all families: *Sp* computed over test data. Stars: *Sp* from links selected with *K*_*link*_(5, *m*) on training sets of different sizes with mutations to alanine only (small stars), 50% (median stars) or 80% (large stars) of data; symbols are reported on panel B. For comparison: results with plmDCA (gray squares), Transformer[53] trained on 80% of data (orange squares). Violet stars: Validation on 36,000 double mutations for RNA binding, and on 3,627 double mutations with large experimental confidence for WW with *K*_*link*_(3, *m*) trained on all single mutations. **D**. Comparison of ranking heuristics based on plm-DCA (p), random (r) (average over 2 realizations) and mutational information (m). *Sp* on train (blue) and test (red) sets for 2-fold cross-validation (average over 10 random partitions, standard deviations are smaller than symbol sizes) for the MutInf procedure at round *r* = 5 are shown and compared to the round *r* = 0 (green).

### Cross-validation on mutations non included in the training set

We then partition the fitness data of the 10 mutational scans under study in two subsets: the training data used by the MutInf procedure to learn the *K*-link model, and the remaining data used to test the model predictions. Cross-validation is repeated for ten random partitions of the test and the train data sets. Figure 3B shows the averages *Sp* and their standard deviations on the 10 partitions, as functions of the relative size of the test set for the *β*lactamase protein ^2^. Panel C provides *Sp* values on the test sets for three value of the training set size and for all the proteins. The Spearman coefficients on the test sets (0.6 *< Sp <* 0.85), are well above the values obtained with the standard plmDCA fully-connected Potts model for all training-set sizes, with the exception of UBOX, for which the improvement with the *K*-link model is modest. The overall average improvement is of about 20% of the Spearman coefficient.

Our results on the 80%-train partition are comparable with the results obtained by the deep-learning software Transformer [53] (black square), with 5 out of 10 families showing similar Spearman coefficients, 3 having worse performances -especially for *β*Glucose, and 2 larger values for *Sp*. Notice that very similar Spearman correlations are obtained with our method with fewer training data, *e*.*g*. 50 − 50% partitions (2-fold cross-validation), but the results for such partitions are not available for Transformer. Figures 3B and C show that the Spearman correlation reaches a plateau when the training set contains more than ∼ 40% of the single mutational data. This robustness with respect to the size of the training set suggests that the set of the most relevant epistatic connections can be inferred, on average, with .4 × 19 ≃ 8 mutations per site only. In other words, when epistatic effects between the mutations carried out by the two sites defining the link are detected for a sizeable fraction of the residues, it is very likely that such effects are also present for other residues. Net improvement with respect to plmDCA is found even when training from a random subset with 5% of the mutations only; this corresponds to train data containing about one mutation on each site only, see first point in Fig. 3B and Suppl. Fig. C.10. A similar result is consistently found in the specific case when the training set includes only the measured fitnesses from the wild-type amino acids to alanine, a frequent and small amino acid (Fig. 3B and C).

### Cross-validation on double mutants

Figure 3 tests the ability of the inferred *K*-link model (trained on single mutations only) to predict about 36, 000 double mutations measured for the RNA Binding domain. We again observe a considerable increase in predictive power with respect to fully-connected models inferred with the standard plmDCA procedure. Figure 3 shows similar results for the WW domain, with a systematic improvement in *Sp* when predicting the mutational effects on the ∼3700 double mutations with higher confidences in the fitness measurements, see also Suppl. Fig. C.10.

### Robustness of the results against the initial ranking of links

We study here how results are affected by the initial ranking of links, by comparing the plmDCA-based ranking (p) with two alternative heuristics:

#### Random ranking (r)

An ordering over all pairs (*i, j*) of sites is drawn uniformly at random.

#### Mutational-scan-based ranking (m)

Compute the Spearman correlation corresponding to the independent-site model, *Sp*_*indep*_. Then, for each one of the *N*_*pairs*_ pairs of sites (*i, j*), compute the variation Δ*Sp*_*i,j*_ between *S*_*indep*_ and the Spearman correlation of the 1-link Potts model with unique connection (*i, j*). Then rank the pairs in decreasing order of Δ*Sp*_*i,j*_.

Figure 3C shows MutIinf selected optimal networks reach similar Spearman correlations when the initial ranking is based on inferred couplings with plmDCA (heuristics *p*) or is purely random (*r*), see also Suppl. Fig. C.12 The MutInf selection procedure alone is therefore able, independently of the initial ranking, to select a good sparse network to support fitness predictions.

Results with the mutational-scan-based heuristics (*m*) show slightly smaller Spearman correlations, see Suppl. Fig. C.12, and a smaller correlation coefficient *R*^2^ for the *β*lactamase, *β*glucose and UBOX (Suppl. Fig. C.13), at the final round of selection.

### Characterization of the optimal epistatic networks

The MutInf selection procedure produces a sparse interaction network supporting the Potts couplings. As shown above, the different heuristics, based on different initial rankings, reaches optimal networks with similar predictive power on mutational effects. However we expect that the selected networks are different, especially for long proteins, as links which are initially low in the ranking are more likely to be pruned due to the increase variance in the network ^3^ (see Suppl. Fig. C.14).

We now investigate how the selected networks inferred with different initial rankings and at different number of rounds compare, and, most importantly, how they are related to the structural and mutational properties of the protein under study. We consider first the case of the RNA binding protein and then extend the results to all proteins and mutational scan under study.

### Core of shared links between equivalent optimal networks

We first compare the interaction networks obtained with MutInf with different initial ranking heuristics. To do so we compute the two optimal networks associated with the *p* and the *m* heuristics, and *n* more networks corresponding to *n* initial rankings produced by the random heuristics *r*. We show the number of common links to these *n* + 2 networks as a function of *n* in Fig. 4A (full curve). This number is much larger than what is expected by chance for the intersection of *n* random networks of the same size (dashed grey line in Fig. 4A, see also Suppl. Appendix A.6). In addition, after an initial decrease, the number of shared links saturates to a large value, indicating the presence of a robust minimal set of links, common to all inferred networks.

**Figure 4.**
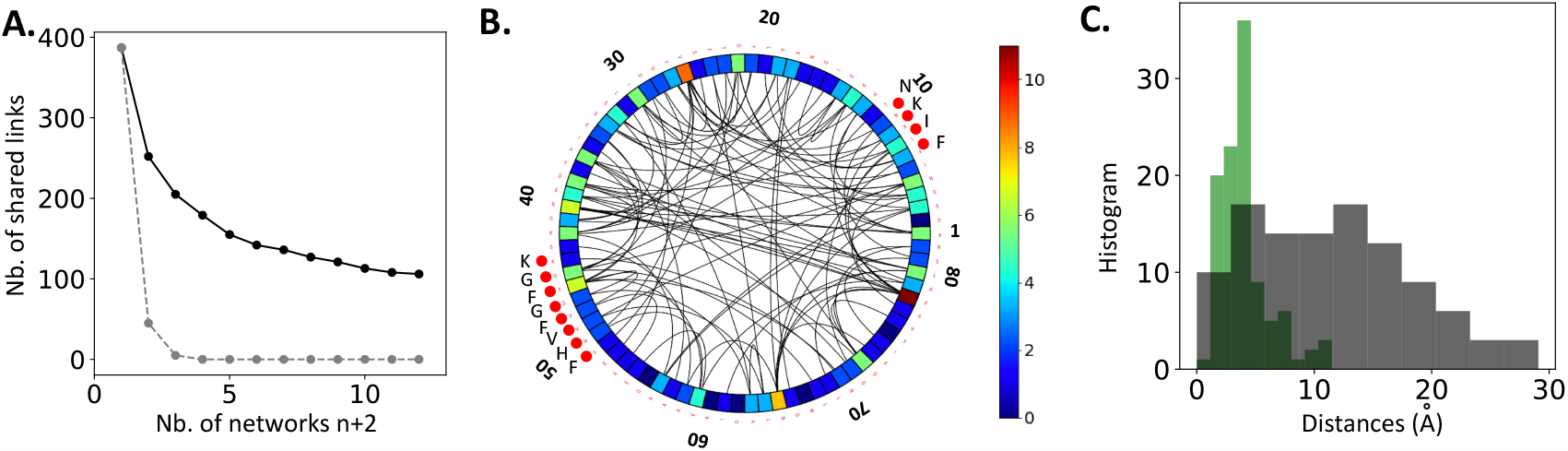
Core network selected by the MutInf pipeline for the RNA Binding protein. **A**. Number of links common to the optimal *K*-link Potts networks corresponding to the *p* and *m* heuristics, and to *n* random initial rankings (full line) vs. total number *n* + 2 of networks. The grey dotted lines shows the outcome for the same networks with randomly permuted links. **B**. Circular plot of the *U*_12_ core network. Colors reflect the site connectivities. The two highly conserved RNP1/RNP2 binding motifs are indicated with red dots and with their amino acids in the wild-type sequence of yeast. **C**. Distribution of carbon-carbon distances between residues in the *K* = 106 pairs of the *U*_12_ core (grey) compared to the ones between residues in the *K* pairs with largest coupling norms according to plmDCA (green). Distances were calculated on the human RNA-bind domain PDBID 1CVJ.

This core subnetwork for *n* = 10 +2, hereafter referred to as *U*_12_, is shown in Fig. 4B. It contains ≃ 30% of links selected by a single optimal network, corresponding to ≃ 3% of the pairs of the sites (see also Suppl. Table B.4). The network connectivity is highly heterogeneous, with some hub-like sites connected to up to 11 other sites, and others to none. These hub sites include the active sites of the protein.

### Distribution of distances between pairs of residues in the selected links

Fig. 4C shows the distribution of distances between the residues connected in *U*_12_, computed from the structure of PABP-Yeast. Distances are broadly distributed around 15 Å, demonstrating that most selected links do not interconnect residues in contact on the protein structure, and have therefore functional rather than structural relevance. In contradistinction the distribution of distances between residues associated with large plmDCA coupling norms, known to be good predictors of contacts [5], is centered around 5 Å.

### Functional interpretation of site connectivities

We have further investigated the relationship between the connectivity of a site and its role in the protein fitness according to the mutational scan. The latter can be quantified in several ways. In Fig. 5A we show the mean fitness on each site, where the average is taken over all tested residues on this site. We also define tolerant and sensitive sites following [41]: a mutation is said viable if the mutant has a fitness larger that the one of the wild-type sequence minus 10% of the fitness interval. Sites having more than 80% viable mutations are said tolerant, while the ones with less than 20% viable mutations are said sensitive (Suppl. Fig. C.11 shows the fitness distributions and the cutoffs for viable mutations on all the mutational scans). Figure 5B shows that most largely connected sites on the *U*_12_ network in Fig. 4 are sensitive or tolerant sites.

**Figure 5.**
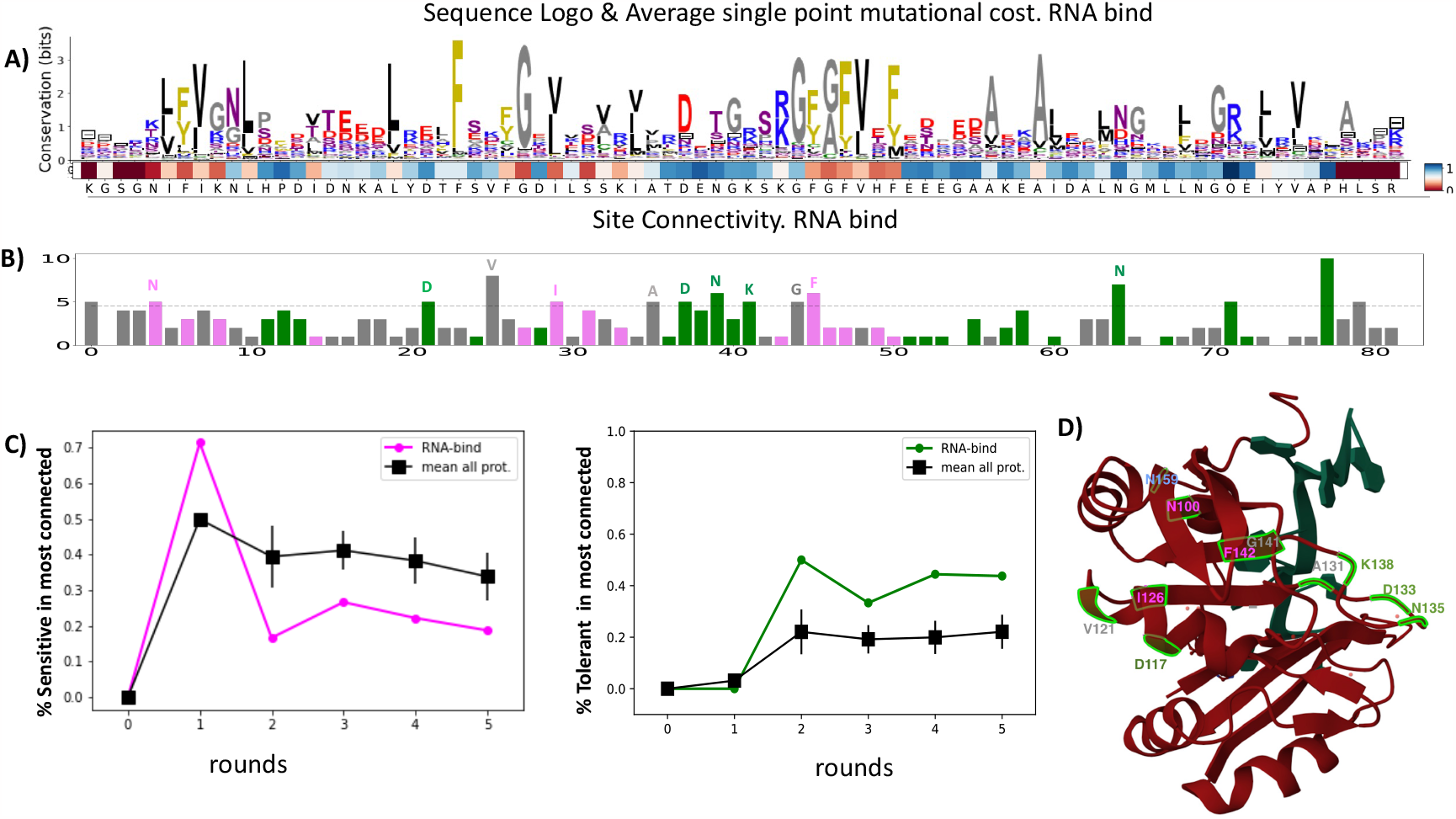
Mutational sensitivity of most connected sites along MutInf iterations. **A**. Sequence Logo of the RNA Binding protein MSA and site average fitnesses, see color code on the right scale. Brown sites are missing mutational data. **B**. Site connectivity in the *U*_12_ network. The dashed line is the threshold used to identify the most connected sites (one standard deviation above the average degree). Magenta/green bars indicate sensitive/tolerant sites. **C**. Fraction of sensitive and tolerant sites among most connected sites, as functions of the round *r* of selection for the *U*_12_ core network of the RNA bind and means over all the mutational scans (error bars are standard errors of the mean) **D**. Most connected sites shown on the PDB Structure 1CVJ of the Poly(A) Binding protein (red) in complex with Polyadenylate RNA (green).

The percentage of sensitive vs. tolerant sites in these most connected ones (having a connectivity larger than the average plus one standard deviation) varies with the round of selection, see Fig. 5C. During the first round of mutationally driven selection^4^, susceptible sites are more likely to support selected links, while, at later rounds, links on sites carrying beneficial mutations are tendentially included. A weaker correlation between connectivity degree and site conservation is observable (see Suppl. Fig. C.16).

In summary the MutInf selection procedure first selects coupling parameters to better predict the effects of detrimental mutations, and, later, to reproduce the effects of beneficial mutations.

### Results for RNA Binding extend to all proteins under study

The main features of the selected networks for the RNA Binding protein above hold for the networks of the other proteins we have studied. As shown in Suppl. Fig. C.15, the dimension of the network shared between *n* different optimal networks first decays with *n* and then reaches a plateau, well above the sharing expected within a null model (see Suppl. Table B.4). Similarly to RNA Binding, selected links do not generally interconnect sites in contacts in the tridimensional structure of the protein; this results also holds for the PDZ (Suppl. Fig C.18) and WW (Suppl. Fig C.19) domains. This feature is consistently shown by the decay of the Frobenius norm of the plmDCA inferred couplings on the selected links with the round of selection, see Suppl. Fig. C.16. Moreover, for all the mutational scans under study, the hub of most connected sites contain the active sites.

Fig. 5 D shows the fraction of sensitive and tolerant sites among the most connected sites. As a general finding the fraction of sensitive sites is maximal at the first round of MutInf selection and then decreases, while the percentage of tolerant sites increases after the first round. Case by case study are presented in Suppl. Fig.C.16.

### Variation of the optimal sparse network with the trait under selection

We now investigate the structure of the selected networks, their differences and similarities for two proteins with multiple mutational scans: the *β*lactamase, for which the effect of single-site mutations on fitness under different antibiotic concentrations have been measured in [21], and Ubiquitin, for which single-mutation effects on the growth rate and on the molecular activation of Ubiquitin upon E1 binding have been measured [41, 42].

### Dependence of the MutInf epistatic network on antibiotic concentration in TEM-1 βlactamase. β-

lactamases are enzymes providing bacteria with resistance against *β*lactam antibiotics. Sensitivity to mutations were shown to largely depends to Ampicillin concentration [21]. Only active sites remain sensitive to mutations at low ampicillin concentrations, at which *β*-lactamase looses its critical effect on the bacteria fitness. Fig. 6A shows the Spearmans between the MutInf predictions and three mutational scans at increasing values of Ampicillin concentration (10, 156 and 2500 *μ*g/mL) measured by [21]. At very low antibiotic concentration the mutational cost predicted from the sequences in the MSA, models inferred from sequence data only, such as the independent-site model and *K*_*link*_(0, *p*) are very bad predictors of the experimental permissiveness to mutations, with *Sp* ≈ 0.2. In other words the sequence data statistics does not reflect such a low selective pressure on the *β*lactamase. With MutInf link selection the inferred models have higher predictive powers on the test data set (*Sp* ≈ 0.6 on test set), but at the price of introducing a complex interaction network, with the many permissive sites having a large degree of connectivity; as the few sensitive ones, which get more and more focused on the active sites as observed in [21], see Fig. 6B,C. This complexity is a hallmark of the disagreement between the rather strong site conservation in the sequence data and the flat fitness landscape at such low concentration. Increasing the antibiotic concentrations, models inferred from sequence data only (no round of MutInf) are more and more predictive (*Sp* ≈ 0.6*/*0.7), see Fig. 6A. With MutInf link selection the inferred models have even higher predictive powers (*Sp* ∼ 0.85) in the presence of a large selective pressure, as already shown in Fig. 3) for the mutational scan of [22]. Correspondingly, the *U*_12_ networks at high concentration are less inter-connected than at low concentration, keeping the large connectivity in the active sites, see Fig. 6C,D. This result supports the idea that natural sequences collected in the MSA are under antibiotic selection.

**Figure 6.**
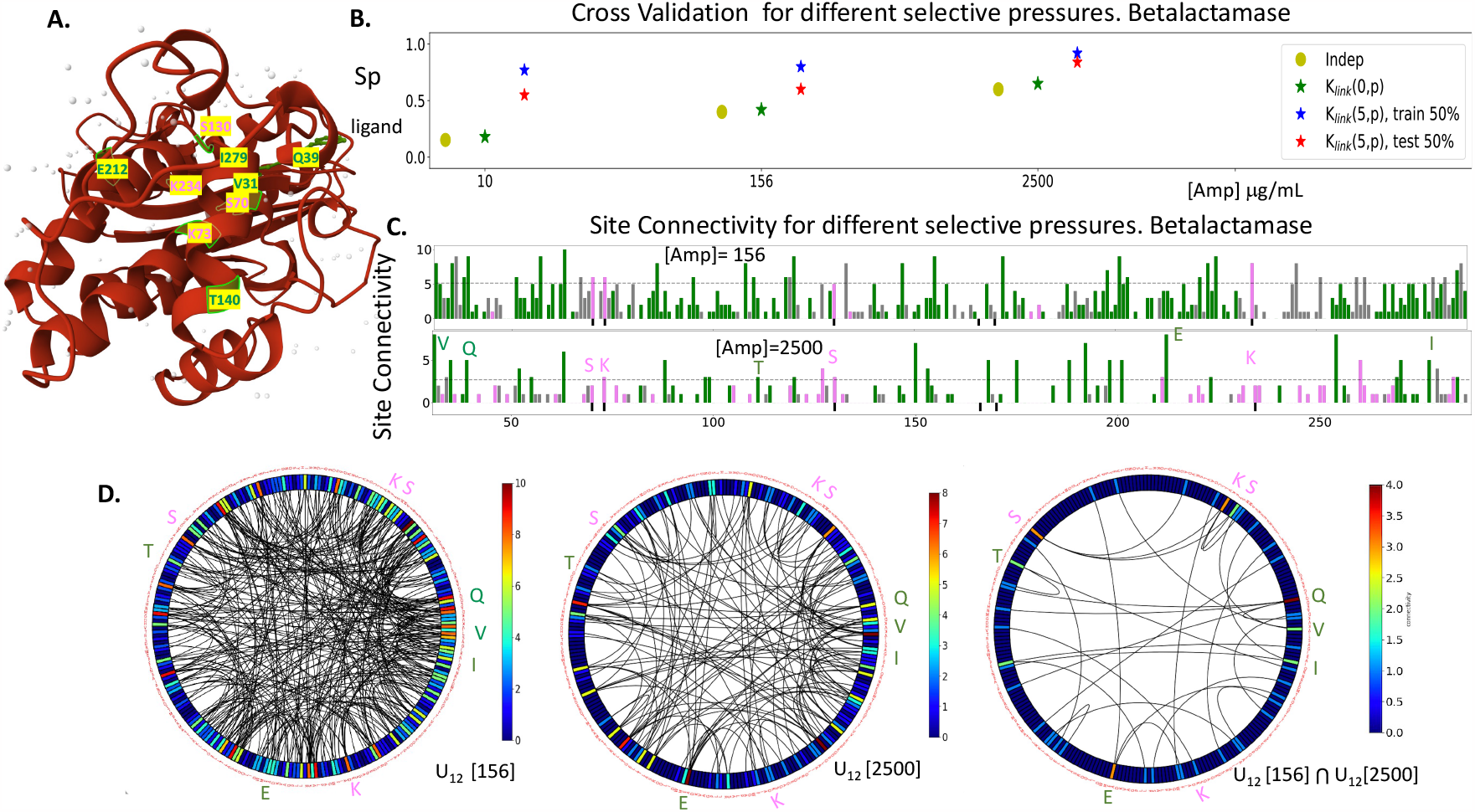
Functional networks of *β*lactamase under variable selective pressures. **A**. Structure of the active state of Tem1 B *β*lactamase from E.Coli (PDB 1AXB) bound to a phosphonate ligand in S70. The most connected residues common to the MutI selected network from mutational scan [21] at 2500 and 156 *μ*g/ml Ampicillin concentrations are displayed: V31, Q39, T140, E212, and the the active sites of the proteins S70, K73 S130 and K234 (amber numbering). Magenta/green labels indicate sensitive/tolerant sites. Similar results are obtained from [22] data, see Suppl. Fig. C.17. **B**. Spearman correlations with the independent-site, *K*_*link*_(0, *p*) and *K*_*link*_(5, *p*) models (train and test set on 10 random 2-fold partitions) on mutational scans at Ampicillin concentrations equal to 10, 156, 2500 *μ*g/ml [21]. **C**. Site connectivity along the sequence for the *U*_12_ networks from the mutational scans at concentrations 156 and 2500 *μ*g/ml. The dashed line is the threshold to identify the most connected sites. Magenta/green bars indicate sensitive/tolerant sites. **D**. From left to right: *U*_12_ networks for concentrations 156 and 2500 *μ*g/ml, with respectively 332 and 135 links, and intersection with 30 links focused around the underlined residues including the catalytic ones.

### Mutational study of RL401 and differences between networks associated with in vivo fitness and E1-activation

Due to its ability to covalently bind to a variety of proteins and complexes, ubiquitin is essential to multiple cellular processes including protein degradation. In [41] the fitness of single-point mutations in *wt* yeast ubiquitin (RL401) gene was estimated through the ratio of the growth rates of the mutants and of the wild-type protein. In [42] the activation of mutants of the ubiquitin protein by E1, the ubiquitin activating enzyme, was directly measured. We compare below the functional networks obtained when applying MutInf to these two mutational datasets. Figure 7A shows the sequence logo in the MSA, and the *U*_12_ site connectivity and network is shown in Fig. 7B,C for the two mutational scans. Key positions required for binding many ubiquitin receptors are, in amber notation, the hydrophobic patch L8, I44, V70, as well as the K48 residue critical, with the C-terminal sites (not included here in the alignment), for covalent linkage. The two networks are both characterized by hubs of largely interconnected sites around the key positions defined above, but their fine structures are different, with only about half of common links between the two *U*_12_ cores (left). In particular the core associated to *E*_1_ activation is more spread along the protein sites, in agreement with the experimental finding that the sensitivity to mutations is strongly correlated with the binding interface with ligands [41] and that the structurally characterized ubiquitin-E1 interface is large and encompasses the interfaces of ubiquitin with most other known binding partners [42].

**Figure 7.**
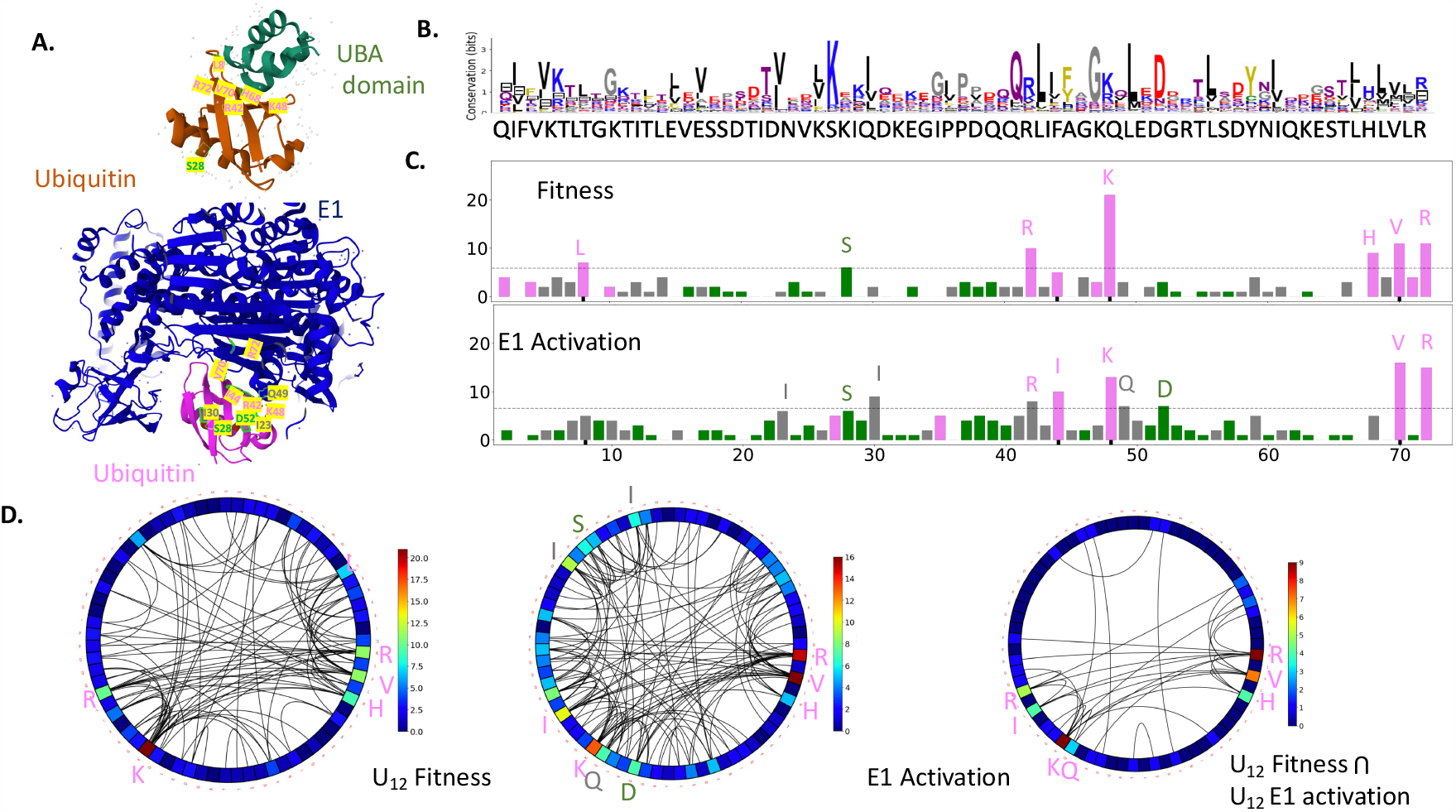
Functional networks corresponding to different fitnesses of Ubiquitin. **A**. Top: Structure of Ubiquitin in complex with the UBA domain from PDB 2OOB. The residues with largest connectivity, L8, S28, R42, K48, H68, V70, and R72 are shown. Bottom: Structure of Ubiquitin in complex with E1, from PDB 3CMM. The residues with largest connectivity, K27, I30,I44, K48, D52, V70, and R72 are shown. **B**. Sequence logo of the MSA. **C**. Site connectivity along the sequence for the *U*_12_ networks from the mutational scans for the fitness and the E1 activation. The dashed line is the threshold to identify the most connected sites. Magenta/green bars indicate sensitive/tolerant sites. **D**. *U*_12_ core networks selected from the two scans data (with, respectively, 90 and 117 links) and their intersection (32 links), focusing around the key residues L8, I44, V70 for binding many ubiquitin receptors, as well as K48, involved in covalent linkage.

## 3. Discussion

In this work, we have proposed a semi-supervised machine learning method to predict mutational fitness costs. It combines the inference of the parameters of a sparse graphical model (*K*-links Potts model) from homologous sequence data with an iterative procedure informed by some mutational data (MutInf) to select few, relevant interactions between residues. This procedure has high predictive power on test data sets, comparable to other state-of-the-art pipelines based on much more complex architectures, while being much more interpretable [53].

The *K*-links Potts model accounts for epistatic effects on the selected links. It has therefore more predictive power than simpler independent site models, which were recently shown to be successful in predicting single mutational effects, when sequence data are accurately chosen [67], phylogenetic information is retained [49], or some mutational data is used [60]. Thanks to its sparsity the *K*-links Potts model offers some advantages with respect to deep neural network models [53]. First it is computationally very fast. It can therefore be applied to large protein sequences and could be extended to a genomic scale. Second, it needs a small amount of training data used in a semi-supervised way, only to guide the MutInf link selection procedure. Third, it selects an interpretable interaction graph specific to the selection pressure applied during the mutational scan. In the following we discuss in further details the three advantages above in relationship to other works, as well as future development of the methods and potential applications.

### Computational advantages of using low–complexity, sparse graphical models

The method introduced here has, informally speaking, low computational complexity. Thanks to the sparsity of the interaction graph, a reduced number of coupling parameters have to be inferred. Moreover our inference algorithm is very fast, being based on a semi-analytical 2-cluster approximation [71, 68, 72]. The latter is exact only for a tree-like structure with no loop in the interaction graph. It would be possible to go beyond the 2-cluster approximation for the loops present in the selected *K*-links graph by using the adaptive cluster expansion at higher orders [71]. More generally other approximate inference schemes could be used for the inference of the field and coupling parameters of the model such as pseudo likelihood, auto-regressive models, Monte-Carlo learning. The pipeline introduced here, due to its low complexity, could be applied not only to predict mutational costs on a single protein but on protein complexes [19, 73] and, more broadly, to the genomic scale to predict viral evolution [27], or the effects of missense mutations in the human genome [46, 47, 74, 75, 76].

### Bias variance trade-off in of low–complexity models

In a related paper [67] we have shown that models for fitness predictions obey a general bias-variance trade-off. The bias in prediction is larger for lower complexity, *e*.*g*. site–independent or sparse models, while the variance of the prediction estimator increases for more complex models due to the limited amount of sequence data compared to the higher number of model-defining parameters [67]. An optimal trade-off between these two opposite effects can be reached by focusing in a controlled way the sequence data around the wild-type protein according to a signal-to-noise criterion [67]. Similarly, the maximal number of links included in our sparse models can here be predicted based on a signal-to-noise cutoff. As a result of this constraint on the maximal number of links, the optimal network obtained with the the MutInf procedure depends on the initial ranking heuristics, even if a stable core of links shared for all the initial ranking heuristics is found. It would be interesting to study how the depth of the alignment can be optimally chosen to improve predictions for the MutInf procedure and how it is related to the flexibility and stability in link selection [67, 77].

### Advantages of using the mutational scan only to select the K links

The MutInf procedure, performs at the state of the art for semi-supervised performances [53], but, similarly to low-N approaches [54, 61], requires a low amount of mutational data, used in our case to select the links. Even when the selection of the links is only guided by tested mutations to alanine, the performances of the procedure improve with respect to the unsupervised counterpart and the fully connected graphical model with plm-DCA[50]

We have used here only single point mutational scans to select links. It would be interesting to include some multiple mutations in the link selection procedure to see how the selected networks change. The iterative pipelind could also incorporate the experimental scan, by testing the multi-loci mutations which are theoretically predicted as the most informative, in a close loop setup between experiments and machine learning modeling. Such a framework is necessary to overcome the impossibility of extensively sample the fitness landscape of a protein given its huge dimensionality. Our approach could also use other bio-informatic pipelines to drive link selection, such as softwares based on structural prediction when predicting mutational effects on the stability or binding interactions of a protein or of a complex [56, 78, 4, 57, 79, 80]. Our pipeline could finally be applied to other inference frameworks to combine specific experiments with information already collected in databases such as the analysis of gene expression data [81].

### Interpretability

The MutInf procedure prunes the covariation information from the MSA through link selection. The selected links unveil both the key sites and the network of interactions related to the functionality experimentally tested. Interestingly the selected links are only marginally related to residues in contact in the tridimensional structure of the protein, contrarily to the DCA couplings [82]. These links could, on the contrary, be related to indirect and effective interactions mediated by the substrate in the functionality under selective pressure, and to false-positive contact predictions in DCA-methods [83]. Selected links are in majority focused on sensitive sites in the first rounds of the MutI-selection, while they also focus on the sites carrying beneficial mutations in latter rounds. They do not reflect only the functionality under investigation in the mutational scan *eg*. the resistance to the antibiotic but also depend the selective pressure used, *eg*. the antibiotic concentration [21].

Natural MSA are not always good sequence data to build a model to predict fitness effects, *e*.*g*. when the wild-type sequence is not well represented in the MSA [67] or when the functionalities experimentally tested are not the driving selective force on the protein evolution family. The mutational scans on *β*lactamase at different concentrations perfectly illustrate this point [21]. At small antibiotic concentration *β*lactamase is not under selective pressure. Models inferred from MSA, which corresponds to sequences naturally selected against antibiotics, have low performances, as shown by the low *Sp* coefficients. As the antibiotic concentration increases models inferred from the MSA have better and better performances. Selecting couplings with the MutInf procedure allows the model to correct this bias and to reach high *Sp* values for all concentrations (Fig. 6A). We hypothesize that, when the *Sp* obtained after MutInf-selection significantly outperforms the initial one, *e*.*g*. as found with BRCA1, Ubiquitin, Ubox, natural conditions are different from the experimentally realized selection.

## 4. Methods

### 4.1. Data

We have applied our pipeline to 10 mutational scans on 9 proteins, see Table 1. The predictability of the mutation-induced fitness effects with Potts models inferred with pseudo-likelihood inference method (Evmutation) was already studied in [50] on the same data sets. The corresponding Spearman correlations are reported in Table 1. We use the multi-sequence alignments and mutational data provided by Evmutation [50] to facilitate comparison, see Supplementary Material therein. Sequences were reweighted to smooth out inhomogeneous sampling effects, see Appendix A.1.

### 4.2. Site entropy and conservation

The site entropy is *S*_*i*_ = − Σ*a p*_*i*_(*a*) log *p*_*i*_(*a*), where *p*_*i*_(*a*) denotes the empirical frequency of amino-acid *a* on site *i* estimated from the sequence data. Site conservation is obtained as *C*_*i*_ = log_2_(21) − *S*_*i*_, and represented by global height of the sequence logo, see Fig. 5A. Sites *i* with entropy *S*_*i*_ *<* 1 are referred to as ‘most conserved’.

### 4.3. Inference of K-links Potts model

We infer the Potts model parameters in the so-called dissensus gauge, see Appendix A.2, in which the fields and couplings attached to the least probable amino acid *d*_*i*_ on each site *i* are set to zero: *h*_*i*_(*d*_*i*_) = 0, *J*_*ij*_(*d*_*i*_, *a*_*j*_) = 0, *J*_*ij*_(*a*_*i*_, *d*_*j*_) = 0, *J*_*ij*_(*d*_*i*_, *d*_*j*_) = 0. We use a color compression approach, in which only the *q*_*i*_ ≤ 21 amino acids + gap symbol present on each site *i* in the sequence data are explicitly modeled [84]. The fields associated to non-observed amino acids *a*_*no*_ are *a priori* set to *h*_*i*_(*a*_*no*_) = − log[10 *B p*_*i*_(*d*_*i*_)], and the corresponding couplings to zero: *J*_*ij*_(*a*_*no*_, *a*_*j*_) = *J*_*ij*_(*a*_*i*_, *a*_*no*_) = 0, see [84]. Here, the *p*_*i*_(*a*_*i*_)^*′*^*s* are the frequencies of the *q*_*i*_ amino acids *a*_*i*_ on site *i*. Our inference procedure for the *K*-link model proceeds as follows. We first compute the values of the fields 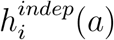they would have in the independent-site model, *i*.*e*. in the absence of any link. The parameters 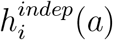 are computed through the numerical minimization of the *L*_2_ norm regularized cross entropy

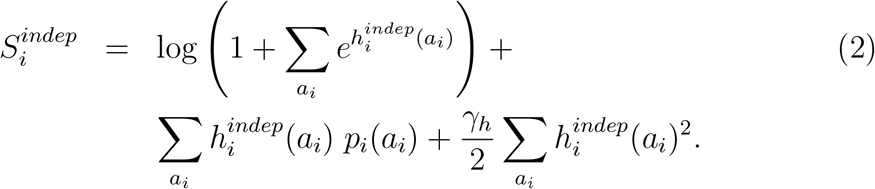

We here choose *γ*_*h*_ = 0.1*/B*, where *B* is the number of sequences. These independent-site values are used to initialize the fields:

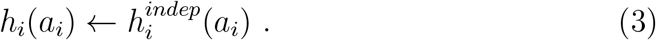

Then, for every link *k* = 1, …, *K* joining the sites *i ≡ i*(*k*), *j ≡ j*(*k*), we compute the couplings *J*_*ij*_(*a*_*i*_, *a*_*j*_) and fields 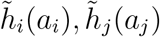through numerical minimization of the regularized cross entropy in the 2-site cluster approximation [71, 68, 69]

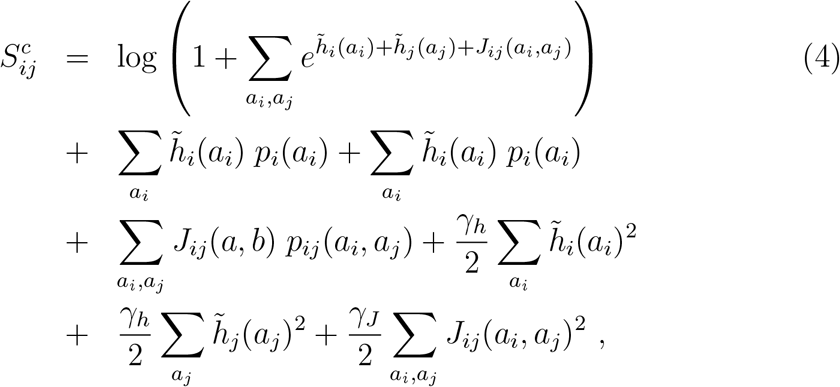

where *γ*_*J*_ = 10*/B*, and the *p*_*ij*_(*a*_*i*_, *a*_*j*_)^*′*^*s* are the joint frequencies of the *q*_*i*_ × *q*_*j*_ pairs of amino acids *a*_*i*_, *a*_*j*_ on sites *i, j*. The inferred coupling matrix is memorized, and the fields are updating according to

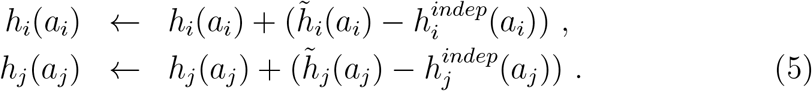

The procedure is repeated over all *K* links. This method relies on the assumption that couplings can be inferred independently of each other, which is generally not true for large values of *K* but seems to hold for the sparse graphs corresponding to the optimal *K*-link networks.

Once the couplings and fields are calculated as described above they are transformed in the wild-type (*wt*) gauge, see Suppl. Appendix A.2. The predicted fitness cost of a single mutation is then simply estimated as

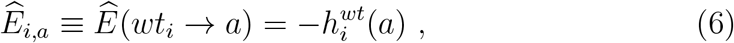

as the field for the wild-type amino acid vanished in the *wt* gauge. The mutational cost corresponding to a 2-point mutation is estimated as

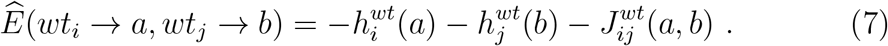

## 4.4. Statistical confidence over single-point mutation predictions

To assess the effect of limited sequence data on the fitness predictors 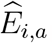 we estimate the variance of the inferred fields in the *wt* gauge, see Eq. 6 and Suppl. Appendix A.2.

In the case of an isolated site (not connected to others) the variance is easy to compute as the fields depend on the frequencies *p*_*i*_(*a*_*i*_) only, with the result

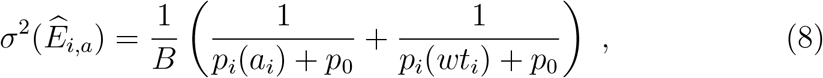

where *p*_0_ = 1*/B* is a pseudo-count used to regularize the expression for unobserved amino acids, *i*.*e*. such that *p*_*i*_(*a*) = 0.

For a site *i* with *K*_*i*_ *>* 0 links to other sites *j*(*k*) with *k* = 1, …, *K*_*i*_, we obtain, according to Eq. 7 the following expression for the variance

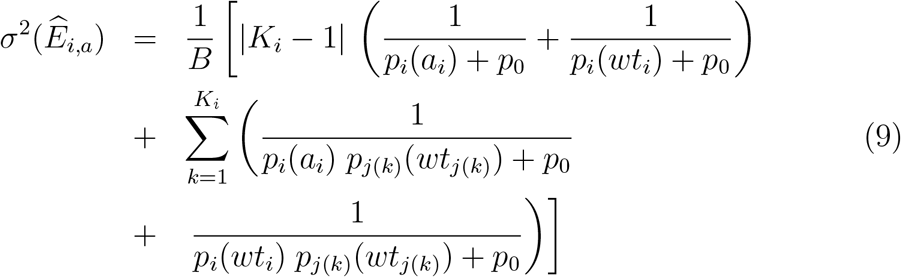

See Suppl. Appendix Appendix A.3 and [69] for the derivation of the above equation. Notice that the approximation used to compute the variance is valid for sparse models only. In the fully connected Potts model inferred by DCA, a strong regularization strength for the coupling parameters of the order of 1*/N* is effective to reduce the contributions to the variance due to the couplings [70].

## 4.5. Criteria for estimating the optimal number of links

The optimal number of links can be guessed based on the following

- Signal-to-noise ratio (SNR) criterion. We compute the average squared mutational variations over all single mutations *wt*_*i*_ → *a*. This quantity, denoted by 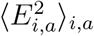 is compared to the statistical standard deviation of the estimators, 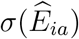, see Eq. 9. We then consider that fitness variations cannot be reliably inferred if the signal-to-noise ratio

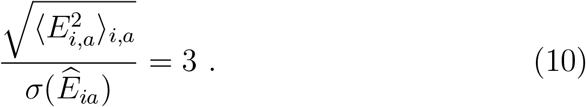

As 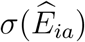 increases with the number of links, see Eq. 9, this arbitrary threshold value effectively captures the number of links the model can include without overfitting the data.
- Robustness-to-noise-addition (RNA) criterion. Alternatively we can add to the predicted mutational cost 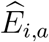 a Gaussian random noise with zero mean and variance *ϵ*^2^, and compute the Spearman correlation (*Sp*) between the predictions of this corrupted model and of the original one as a function of *ϵ*^2^, see Suppl. Fig. C.9. *Sp* abruptly drops for large *ϵ*^2^. We define the maximal size of the network giving the data the one at which value of the standard deviation *ϵ*^2^ gives a Spearman Coefficient decrease to 90% of its initial value (see Fig. C.9).

These two criteria give similar and excellent predictions for the maximal number of links *K*^*∗*^, with very small loss in terms of Spearman correlation with respect to the optimal network determined by scanning all possible *K*, Table B.2. Notice that the criteria do not take into account the model bias. A complete analysis of the bias-variance trade-off for fitness predictions with graphical models is given in [67].

## 4.6. Details about the MutInf selection procedure

### Pruning

At each round of MutInf links with negative Spearman contributions are removed. Keeping these links makes the procedure slower, the final network bigger, and performance worse, see Suppl. Fig. C.12

### Optimal network

The optimal number *K*^*∗*^ of links is the minimal *K*, such that the *K*-link Potts model predictions maximizes the Spearman correlation with the experimental fitness measurements (within relative accuracy of 0.5%). This choice corresponds to the minimal network with large Spearmann. The difference with the network corresponding to the maximum Sp is only relevant at the round 1 which dispalys,around the maximal Sp, a large plateau of Spearman values as a function of the network size *K*, see Fig.2.

### Trimming the initial list of links for long proteins

For long proteins *N*_*pairs*_ is very large, and the links with low ranks in the initial lists have small *Sp* contributions C.14 as the variance increases with the number of links added in the network (C.8). We have verified on *β*Lactamase and *β*Glucose that setting a cut-off *K*_*max*_ = 6000 on the initial list length, performances are weakly degraded with respect to the outcome of MutInf on the train set with the full list, see C.12, and are practically unchanged on the test set for the 0.5-0.5. With this simple procedure the computational time is reduced by a factor ∼200 (Intel^®^ Xeon(R) CPU E5-2690 v4 @ 2.60GHz x 56 (20 cores used)). To spare time in the Crossvalidation procedure for the *β*Glucose we have then used such cut-off.

## Acknowledgments

The authors are grateful to F. Rizzato for her contribution in data processing, and to R. Ranganathan for suggesting the analysis of his *β*lactamase scan. This work was supported by Grant No. ANR-19 Decrypted CE30-0021-01.

## Appendix A. Supplementary Methods

### Appendix A.1. Sequence data: reweighting

The sampling of biological sequences in the MSA is biased by the phylogenetic history of the proteins and by the human selection of sequenced species. To reduce this effect, we decrease the statistical weight of sequences having many similar ones in the MSA. More precisely, the weight of each sequence is defined as the inverse number of sequences within Hamming distance *d*_*H*_ *< xL*, with *x* = 0.2 [85].

### Appendix A.2. Change of gauge of fields and couplings parameters from the dissensus to witd type

Couplings and fields are easily transformed into the *wt* gauge as follows:

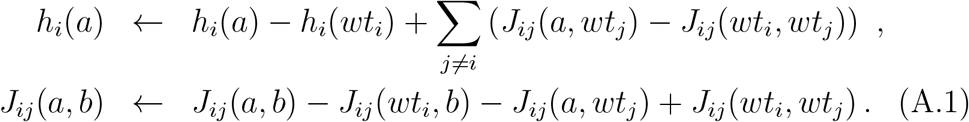

### Appendix A.3. Estimation of the variance on the predicted cost of mutation

We introduce a small pseudo-count 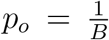 in the expresssions of the fields and couplings. Applying the gauge transform in Eq. (A.1), we obtain

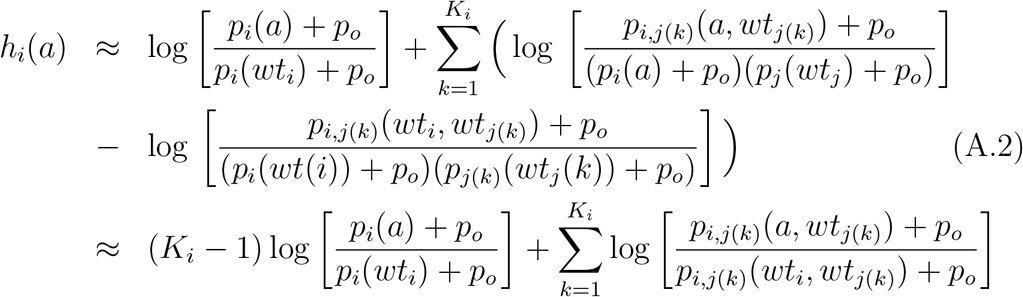

Next we approximate the variance of log(*f* + *p*_*o*_) with 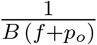, where *f* is a frequency, 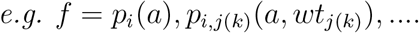 Finally we sum up the variances of the different log terms in Eq. (A.2).

### Appendix A.4. plmDCA used for initial ranking of links

To obtain the initial ranking of links in the plmDCA heuristic we have used the plmDCA algorithm implemented in [82] and adapted to compressed Potts model inference in which amino acids not present in the MSA are not modeled [84]. We have also used reweigthing parameter 0.2, see Appendix Appendix A.1, and large regularization *λ*_*j*_ = *λ*_*h*_ = 0.01 on couplings and fields parameters. Before computing the Frobenius norm *F*_*ij*_, the couplings are transformed in consensus gauge Appendix A.2 and corrected with Average Product Correction: 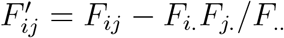 where *F*_*k*._ = Σ _*k*_ *F*_*i,k*_ [86].

### Appendix A.5. plmDCA used to predict fitness costs

The reference plmDCA fitness predictions are obtained as before with the compressed Potts model inference and reweighting parameters *w* = 0.2, but with the a regularization strength for the couplings decreasing with the number *N* of sites in the protein *λ*_*J*_ = *N/B*_*eff*_ *λ*_*h*_ = 1*/*(10*B*_*eff*_), where *B*_*eff*_ is the effective number of sequences after the reweighting procedure, as in [50]. Results with this plmDCA procedure (dark gray squares in Fig. 2) are comparable to the ones given in [50] and listed in Tab.1.

### Appendix A.6. Null model for the number of shared links as a function of the number of compared networks

We consider a network with a maximal number of links *K*_*tot*_ = *N* (*N* − 1)*/*2, the probability to have a common link in *n*_*r*_ random networks of siz 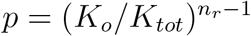. The probability to have *K*_*c*_ common links follows then a binomial distribution with probability *p* and number of extractions *K*_*o*_, the average value (orange curve in Fig. C.15) is *m* = *p* × *K*_*o*_ and the p-value for *K*_*o*_ is computed as the sum of the binomial probabilities for *k* running from *K*_*c*_ to *K*_*o*_. For large *n*_*r*_ and small *p*, the p-value can be estimated from the value of the Poisson probability for *K*_*c*_, using the Stirling approximation. Table B.4 gives the p-values for the fraction of common links for the *U*_12_ core optimal networks:

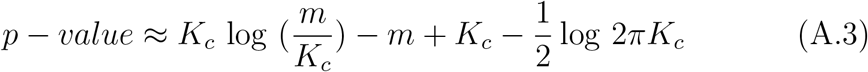

## Appendix B Supplementary Tables

**Table B.2:** Estimates of the optimal number of links. The table gives for each protein the optimal number of links *K*^*∗*^ and the corresponding maximal Spearman *Sp*^*∗*^ for *K*_*link*_(0, *p*) and *K*_*link*_(2, *p*), the relative Spearman loss Δ*S* = *Sp/Sp*^*∗*^ 1 (within 1% accuracy) when estimating the optimal *K* with the SNR and RNA criteria.

| Protein | Round 0 |  |  |  | Round 2 |  |  |  |
| --- | --- | --- | --- | --- | --- | --- | --- | --- |
| | $K^*$ | $Sp^*$ | $\Delta Sp$<br>SNR | $\Delta Sp$<br>RNA | $K^*$ | $Sp^*$ | $\Delta Sp$<br>SNR | $\Delta Sp$<br>RNA |
| βlacta | 350 | 0.66 | 0.01 | 0.03 | 385 | 0.83 | 0 | 0 |
| βgluco | 780 | 0.57 | 0.01 | 0.02 | 1060 | 0.81 | 0 | 0 |
| BRCA1 | 34 | 0.5 | 0.02 | 0.02 | 353 | 0.74 | 0.01 | 0 |
| DNAbind | 386 | 0.59 | 0.14 | 0.08 | 392 | 0.75 | 0.01 | 0.01 |
| PDZ | 571 | 0.56 | 0.06 | 0.05 | 675 | 0.82 | 0.02 | 0.03 |
| Ubi ftn | 97 | 0.5 | 0.04 | 0.09 | 303 | 0.83 | 0 | 0 |
| Ubi E1 | 28 | 0.5 | 0.04 | 0.11 | 282 | 0.78 | 0.01 | 0 |
| RNA-bind | 580 | 0.63 | 0.02 | 0 | 556 | 0.86 | 0 | 0 |
| UBOX | 144 | 0.51 | 0.06 | 0.02 | 366 | 0.71 | 0 | 0.03 |
| WW | 19 | 0.63 | 0.04 | 0.03 | 106 | 0.77 | 0.01 | 0 |

**Table B.3:** Degree of sparsity of the optimal network at the last round of MutI. Proteins are ordrered by their length. Columns, from left to right: Protein name and length; Optimal number of links at round 5 for the plmDCA initial ranking (other heuristics give similar values); Degree of sparsity of the network; Average connectivity degree per site. Number of sequence data in the MSA (See Table 1).

| Prot | N | $K_{opt}(r=5)$ | $K_{opt}/K_{tot}$ | $K_{opt}/N$ | B |
| --- | --- | --- | --- | --- | --- |
| WW | 30 | 87 | 0.2 | 2.9 | 8251 |
| DNAbind | 63 | 333 | 0.18 | 5.6 | 11278 |
| Ubifit | 71 | 264 | 0.1 | 3.6 | 10023 |
| UbiE1 | 71 | 258 | 0.1 | 3.6 | 10023 |
| BRCA1 | 75 | 283 | 0.1 | 3.8 | 14448 |
| UBOX | 76 | 229 | 0.08 | 3 | 14448 |
| RNAbind | 82 | 387 | 0.1 | 4.7 | 70779 |
| PDZ | 84 | 526 | 0.15 | 6.2 | 24795 |
| Blactamase | 253 | 902 | 0.02 | 3.6 | 7620 |
| Bgluco | 441 | 3131 | 0.03 | 7 | 26606 |

**Table B.4:** Fraction of common links in the U12 core network. Proteins are ordrered by their length. Columns, from left to right: Protein name and length; Number of links at round 5 in the *U*_12_ network; Ratio of this number of links and of the optimal network size. The p-value for the *U*_12_ network computed from a random-sharing null model is smaller than 10^−300^ for all proteins.

| Prot | N | $K_{u12}$ (r=5) | $K_{u12}/K_{opt}$ (r=5) |
| --- | --- | --- | --- |
| WW | 30 | 46 | 0.52 |
| DNAbind | 63 | 102 | 0.3 |
| Ubi ftn | 71 | 90 | 0.34 |
| Ubi E1 | 71 | 117 | 0.45 |
| BRCA1 | 75 | 91 | 0.32 |
| UBOX | 76 | 38 | 0.17 |
| RNAbind | 82 | 106 | 0.27 |
| PDZ | 84 | 198 | 0.38 |
| βlacta | 253 | 132 | 0.14 |
| βgluco | 441 | 284 | 0.10 |

## Appendix C Supplementary Figures

**Figure C.8:**
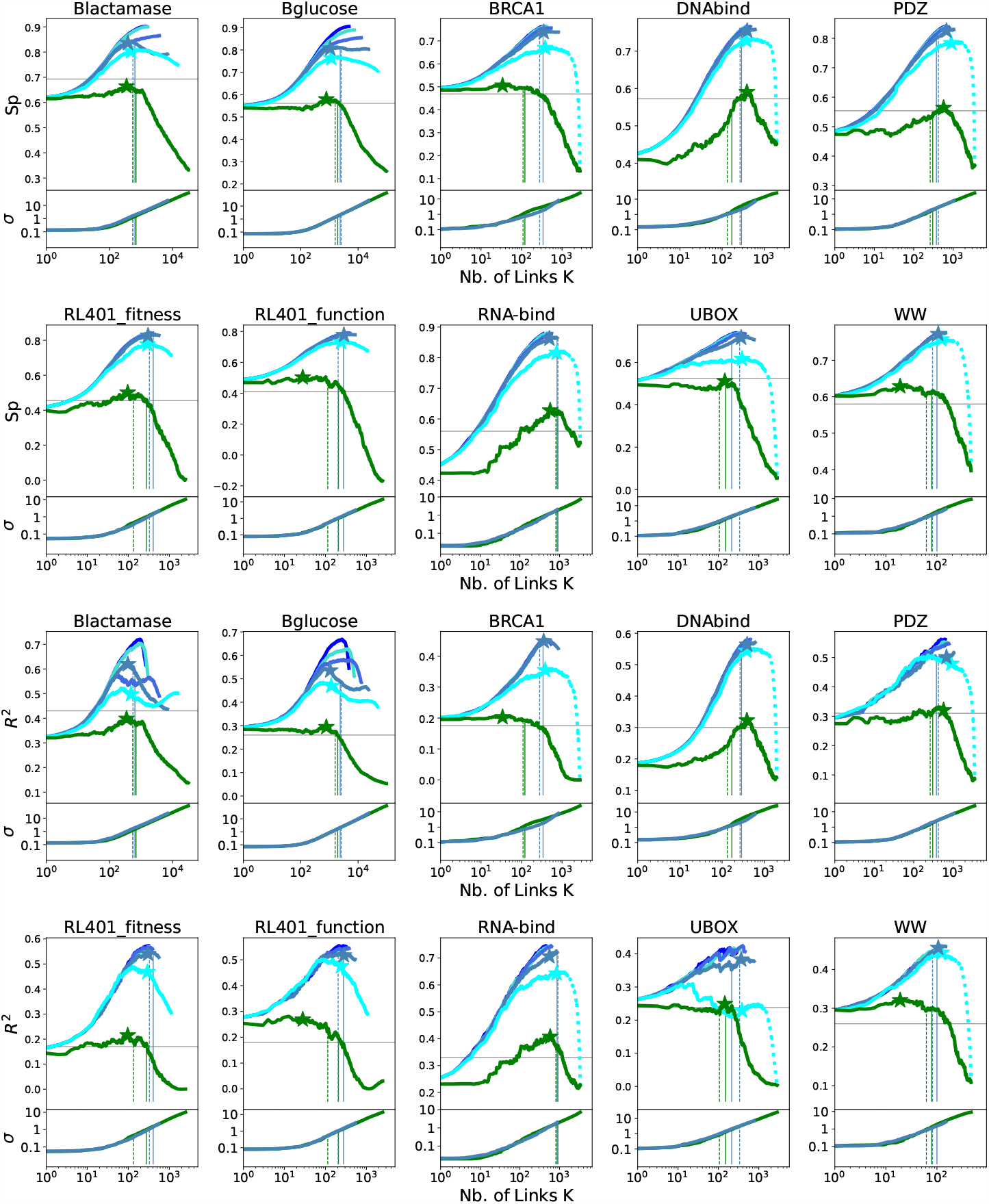
*Sp* and *R*^2^ as a function of the number of links K added for the *K*_*link*_ (p) Potts model for the plmDCA ranking heuristic, for the round 0 (green) followed by 5 rounds of MutI selection (cyan, steelblue,royalblue,turquoise,blue), for all proteins and mutational scans under study (similar to main text Fig. 2 for RNA-bind). Longer is the protein (see Tab.1), larger is generally the number of rounds necessary to obtain convergence in the Spearman: *β*lactamase and *β*glucose converge at round 4, PDZ and UBOX and RNA-bind at round 3 BRCA1, RL401 at round 2 and WW at round 1. For BRCA1, DNAbind, PDZ, RNAbind, UBOX, WW the continuation of the *Sp* vs K on the pruned links is shown with a dotted lined also. The optimal network size *K*^*∗*^ for the first 3 rounds are shown by stars (Sp values given in Fig. 2). Bottom: Variance as a function of *K* for the different models. Dashed and full lines: cutoff network size according to respectively the signal to noise and Spearman robustness to noise criteria Fig. C.9 is predictive of the maximal network size. Gray horizontal lines Plm *S*_*p*_ and *R*^2^.

**Figure C.9:**
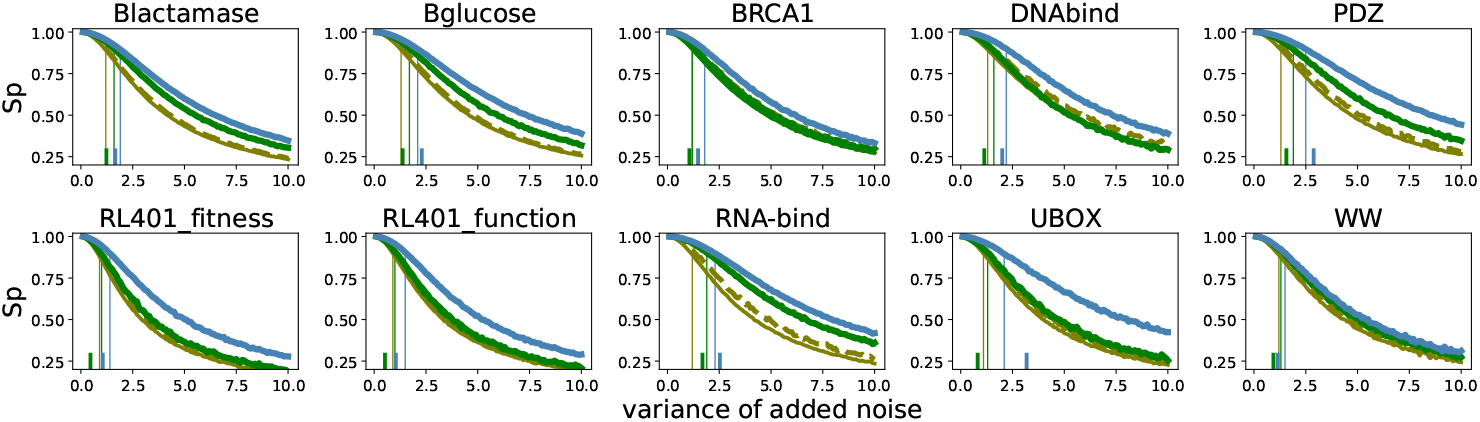
Deterioration of Spearman with a Gaussian random noise added to the predicted cost of mutations as a function of the variance of the noise for all the proteins. Spearman between noise-perturbed predictions and the original ones (full line) and between the noise-perturbed predictions and the data (dashed lines) behaves similarly. Olive green: independent model, green: K_*links*_ (0,p), blue: K_*links*_ (2,p). Vertical full lines determines the cut-off variance at which *Sp* = 0.9. The variance cut-off determined by the signal to noise criterium (used in Fig.2:B), shown by small vertical lines behaves similarly.

**Figure C.10:**
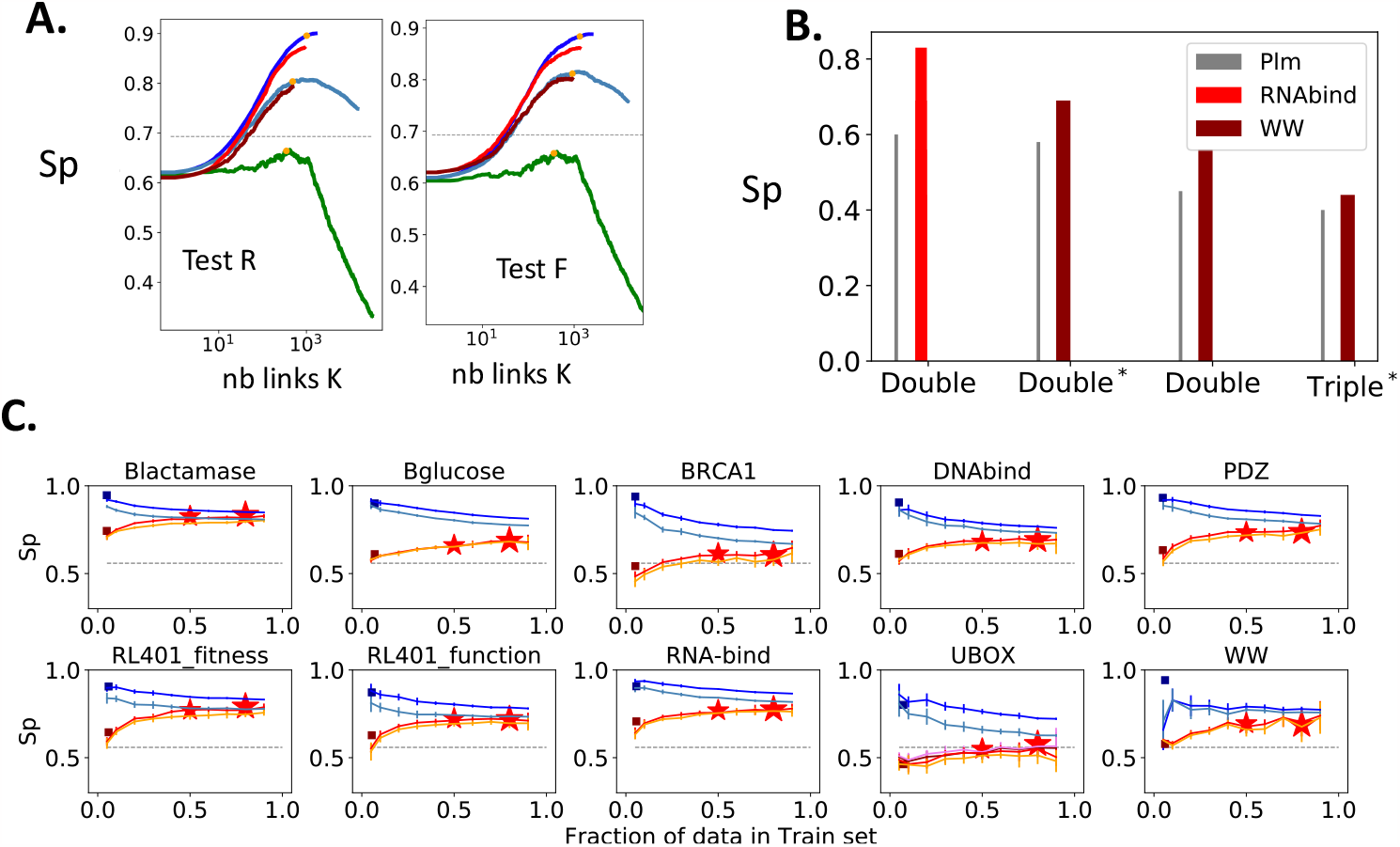
**A**. Cross validation for *β* lactamase on two different mutational scans. *Sp* as a function of the number of links added to the network. Left/right: train/test: Firnberg et al [22], test/train: Stiffler et al [21]: Green, steelblue, blue: MutI (p) Rounds 0,1,4 Sp vs K on train sets. Marron, Violet, Red: Sp on test sets from selected links on train. Yellow dot indicates the network size selected on the train set. **B**. CrossValidation on RNABind Double mutations (36522 data points) and WW Double and Triple mutations (whole set of 6918 data and the more reliable set of 3627 data, for double and 4578 for triple mutations indicated by star), compared to plm-DCA performances. **C**. Spearman Correlation coefficients as a function of the percentage of data used in the train test, for the train test (light blue: K_*link*_(1,p), dark blue: K_*link*_(2,p) and the train set (Orange: K_*link*_(1,p), Red: K_*link*_(2,p) For Ubox 4 rounds of MutI are shown. Stars indicate crossvalidation from selected K_*link*_(5,p) (see Fig.3. Main Text).

**Figure C.11:**
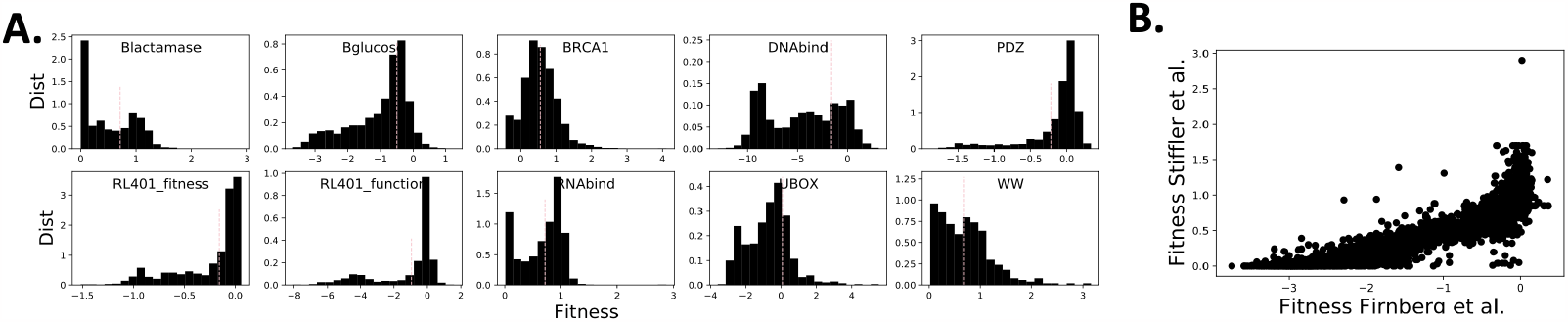
**A)** Distribution of fitness changes in the single mutational scans under study. The vertical bars indicate the cutoff used in the definition of tolerated mutations. **B)** Scatter of fitness values in the two independent single site mutational scan on the *β*lactamase by Stiffler et al. [21] versus Firnberg et al. [22]. The points correspond to the 4591 mutations tested in both scans (among a total of respectively 4788 [21] and 4610 [22]). The Spearman Coefficient between the two data sets is 0.94.

**Figure C.12:**
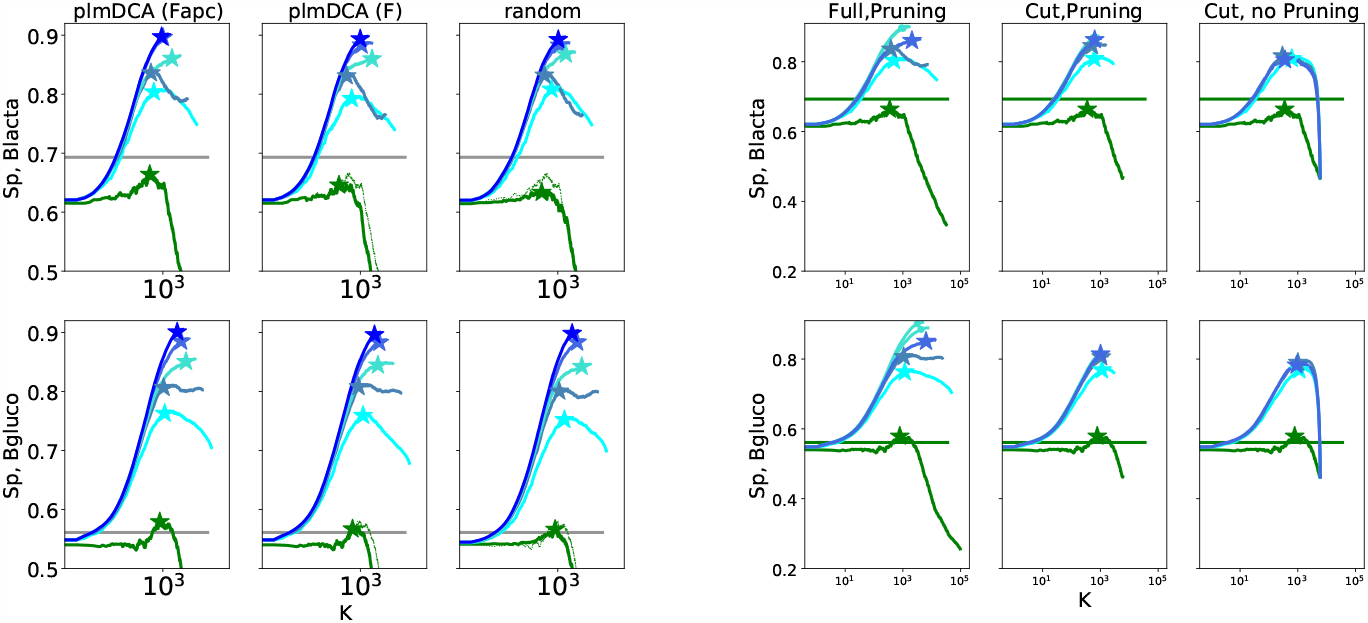
Left: Comparison of heuristics for initial links-ranking for the two longest proteins under study: *β*-Lactamase (top) and *β* Glucose (Bottom). Three heuristics are compared: two obtained from the plmDCA couplings: i) ranking according to the standard Frobenius norm of the couplings with APC corrections *F*_*AP C*_ Appendix A.4 (left) ii) ranking according to the Frobenius norms without APC corrections *F* (center), iii) random initial renking. While the maximal Spearman at round 0 (green) is slightly larger for the plmDCA with *F*_*AP C*_ initial rank (shown in dotted green for comparison) in the following rounds of mutI-procedure the Sp is insensitive to the initial ranging heuristic. A similar behavior is obtained for the other proteins. Right: Comparison on different selection procedures in the successive rounds of MutI selection for the two longest proteins studied,*β*Lactamase (top) and *β*Glucose (Bottom). Three procedures are compared: the standard procedure (starting from the full list of links and pruning links with negative Sp contribution at each round) (left), the same pruning procedure but starting on a reduced list of the first-ranked 6000 links according to *F*_*AP C*_ (center), the procedure consisting in starting from a reduced list of 6000 links but keeping all links in different round of selection, changing their ranking, without removing the ones with negative Spearman contributions (right). This procedure is slower than the standard one because the network does not decrease in size over the round of selections. The full pruning procedure reaches the maximal Sp, the cut-pruning procedure has slightly smaller Sp while the initial cut-non pruning procedure converge faster to a small value of the Sp. Similar results are obtained for the shorter proteins, without the cut in the initial list.

**Figure C.13:**
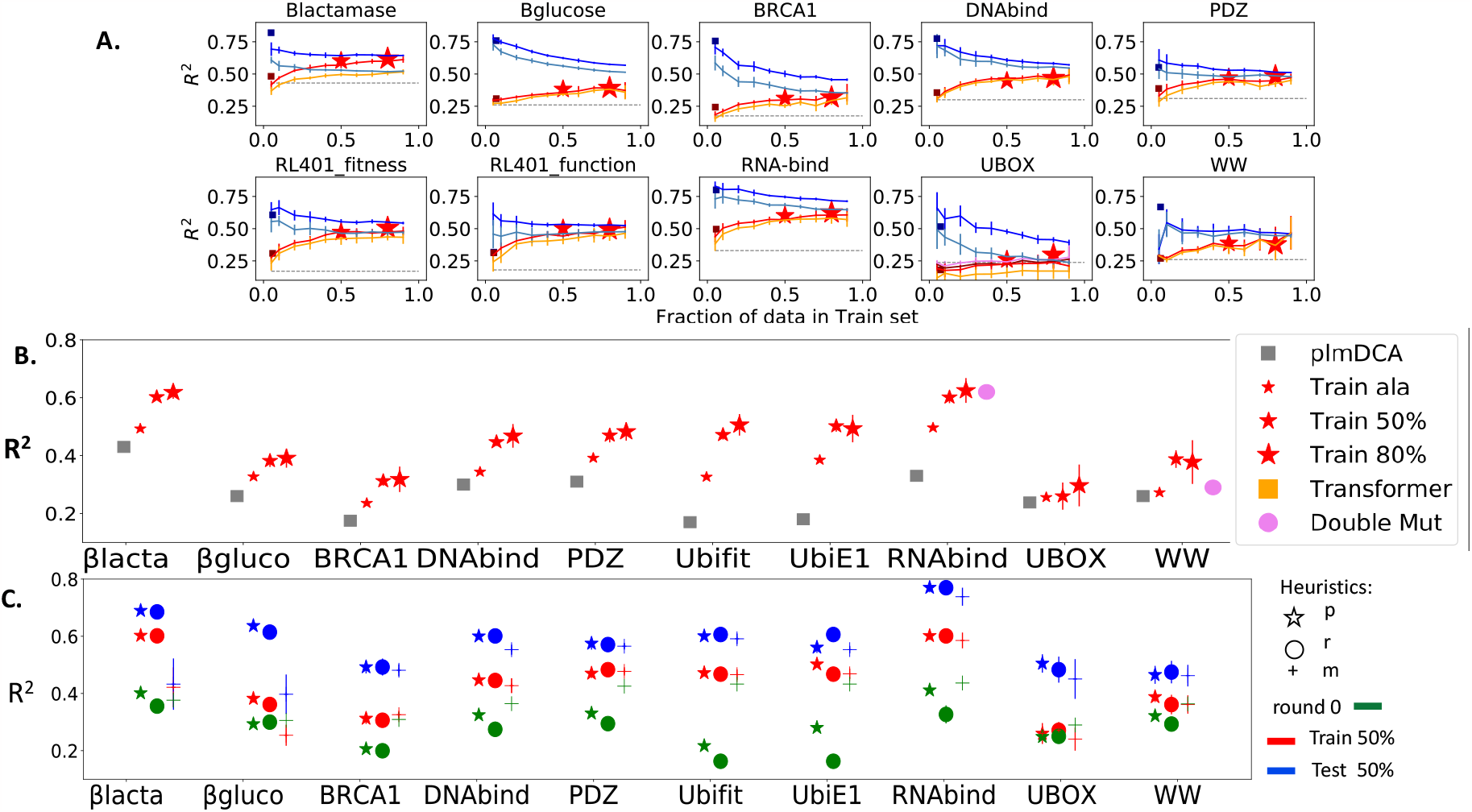
**A**. *R*^2^ Correlation Coefficient as a function of the percentage of data used in the train, for the train test (light blue: K_*link*_(1,p), dark blue: K_*link*_(2,p)) and the test set (Orange : K_*link*_(1,p), Red: K_*link*_(2,p)). Stars indicate crossvalidation from selected K_*link*_(5,p) network (see Fig.3. Main Text). **B**. *R*^2^ for different partition of the Cross-Validation train and test sets. Comparison of performances with Transformer and plmDCA. **C**. *R*^2^ on the 50%-50% partition for the train-test data sets in the CrossValidation procedure for all the heuristics (p,r,m), at round 5 of MutInf compared to the round 0.

**Figure C.14:**
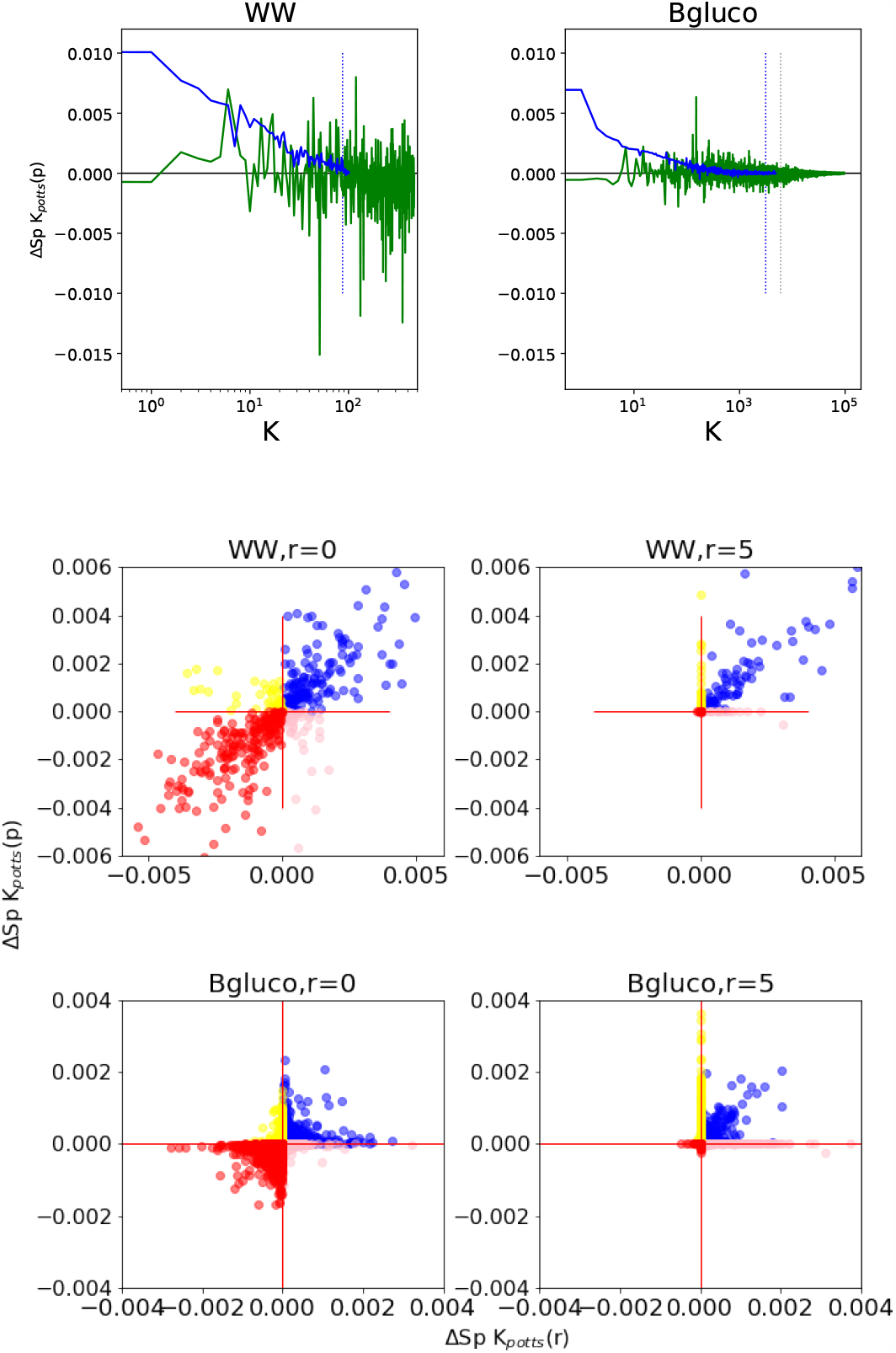
Top: Δ Sp contribution as a function of the number of added links for the plmDCA ranking at round 0 (green) and round 5 (blue) for the WW and *β*Glucose mutational scans. At round 0 positive and negative contributions are present, with an amplitude decreasing with *K* for the longer *β*Glucose; at the round 5 contributions are mostly positive and almost perfectly ranked in decreasing order. Blue dotted lines: the optimal size for the network at round 5, gray dotted line: cutoff to K=6000 used for the crossvalidation for long proteins. Bottom: Scatter of the Spearman contribution Δ*S*_*p*_ obtained at round 0 and round 5 before the re-ranking, from plmDCA (p) initial ranking versus a random (r) initial ranking. Top: WW, Bottom: *β*Glucose. Blue/Red:Δ*S*_*p*_ *>* 0 / respectively *<* 0 for both the p and r ranking, yellow/pink Δ*S*_*p*_ *>* 0 for the p, negative for r ranking or vice-versa. For both proteins at round 0 both positive and negative Δ*S*_*p*_, are present, while at round 5 the Δ*S*_*p*_ are in large majority positive. For the small protein WW the Δ*S*_*p*_ *>* 0 are well correlated and over 208/98 (at round 0/5 respectively) links having positive *Sp* for (p) 162/76, are shared with r, while for large protein the sign-correlation is large but many large contribution with one initial ranking heuristic can be small or zero with other initial rankings. For the *β*Glucose over 34039/2984 links (at round 0/5 respectively) with positive Δ*Sp* for (p) 27239/1322 (80/45%) are shared with (r), this sharing is compatible to the one for 2 optimal network given by *n* = 2 in fig C.15.

**Figure C.15:**
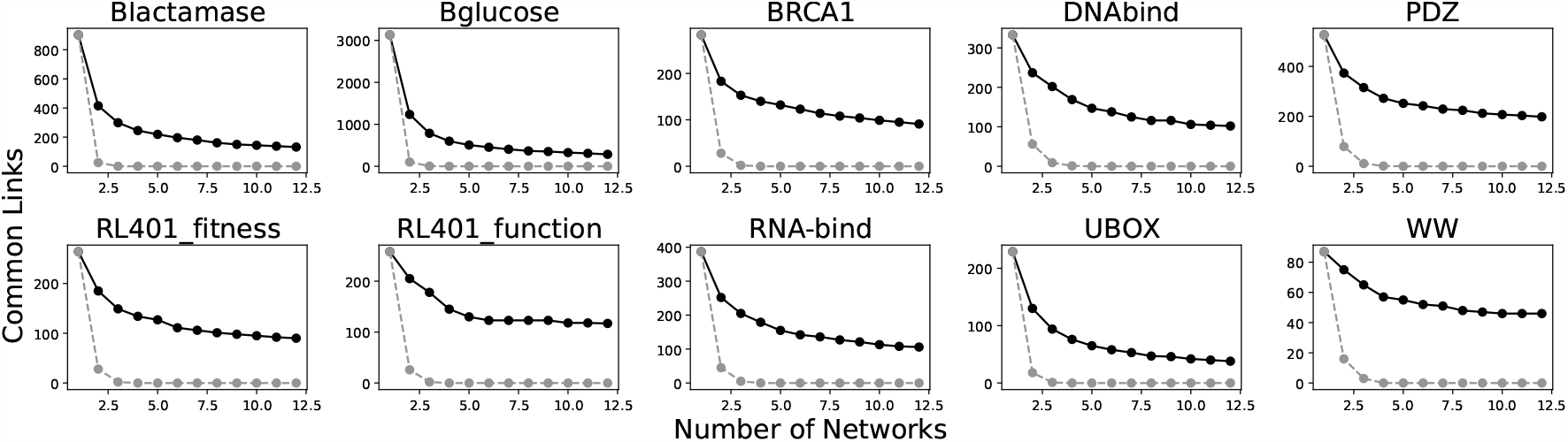
U12 networks for all the mutational scans at round r=5. Full black lines: Number of common links in selected networks, differing by initial sorting of links: the *p* and the *m* heuristics followed by n=10 random initial ranks. The initial number of links is the one for *p* heuristic. Dashed gray line: Null model for random selection. Notice that the number of common links in the U12 are atypical already at the first round of selection.

**Figure C.16:**
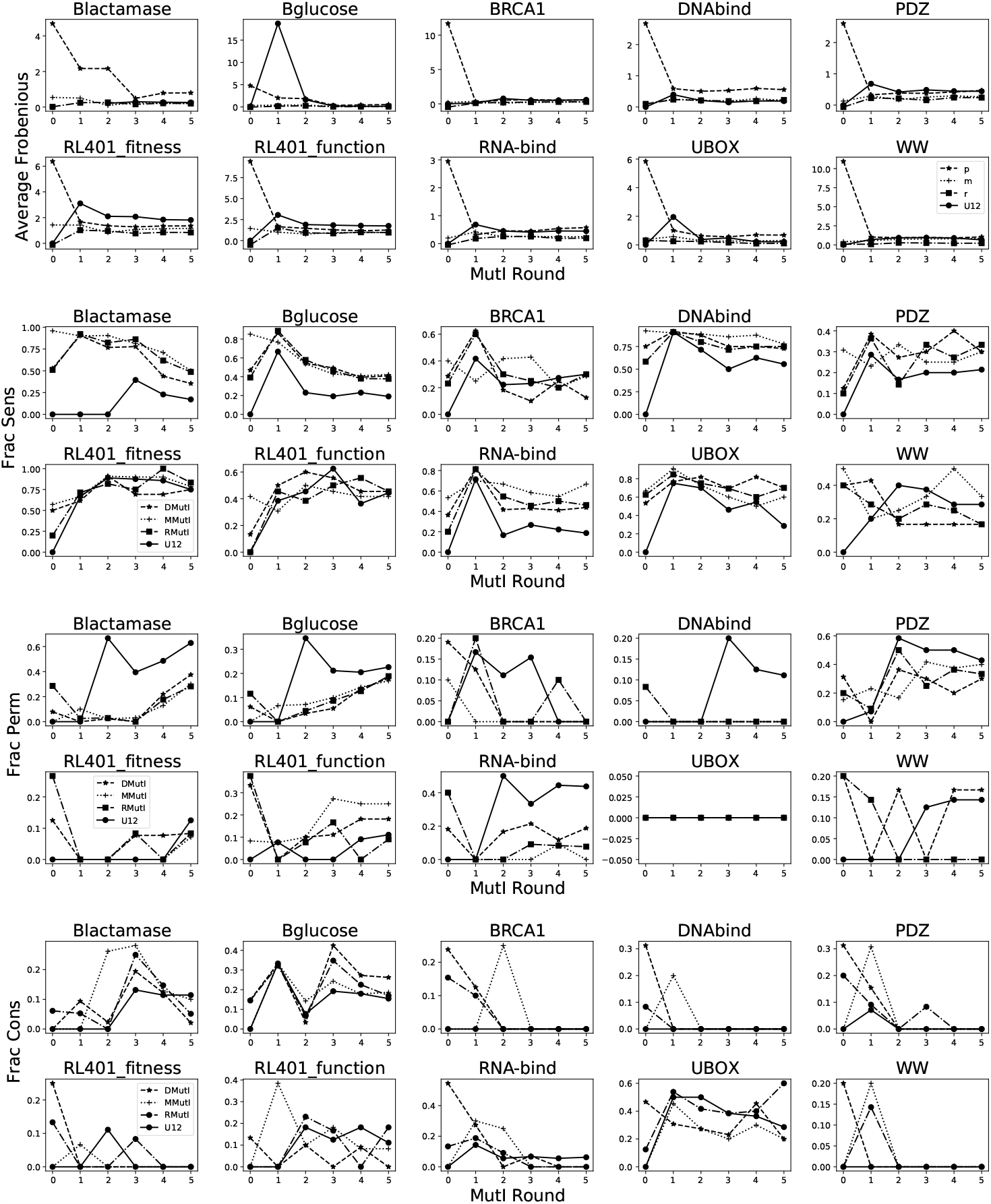
Top: The average Frobenius norm of selected links (computed from the stanbdard plmDCA inference) for different initial ranking heuristics and different rounds decreases due to the MutInf selection starting from from the plmDCA initial lists (m,0), were the Frobenius norm is the ranking criterion. Middle-top: Fraction of most sensitive sites in most connected ones for different selection procedure and different rounds and all the studied proteins. Middle-bottom: Fraction of most permissive sites in most connected ones for different selection procedure and different rounds and all the studied proteins. Bottom: Fraction of most conserved sites in most connected ones for different selection procedure and different rounds and all the studied proteins

**Figure C.17:**
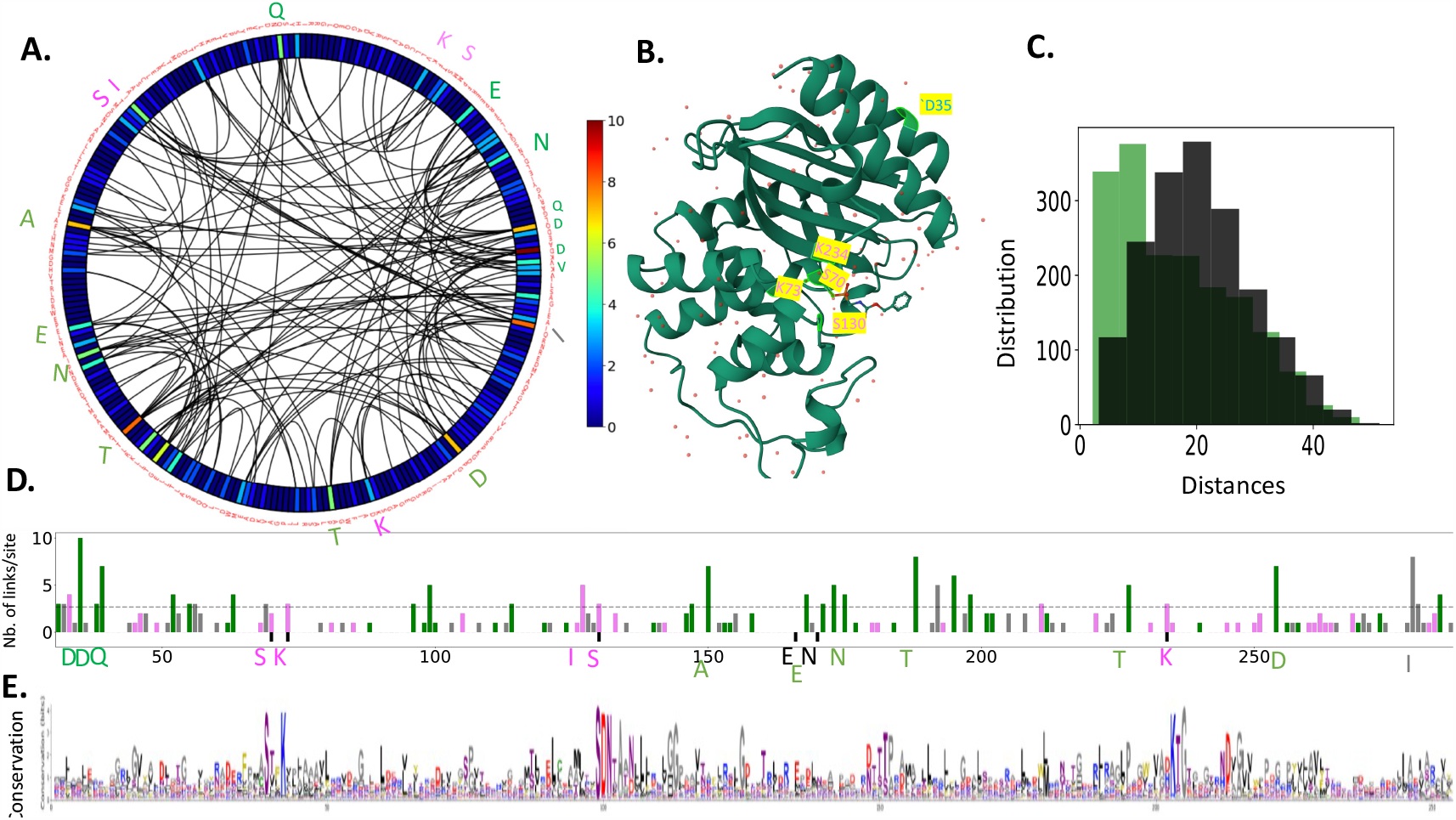
*β*lactamase Structural and functional Couplings: **A**.: U12 network. The site connectivity is indicated by color-code. **B**.. Structure of the active state of Tem1 *β*-lactamase from ecoli (PDB Entry 1AXB) bound to a phosphonate ligand in S70, the 5 most connected residues in the MutI selected network from mutational scan [22] an d underlined: D35 and the the active sites of the proteins S70, K73 S130 and K234 (amber numbering **C**. Histogram of distances for the pairs connected in the U12 network, compared with the histogram of distances for the same number of pairs with largest Frobenius Norm with APC correction. Similar histograms are obtained for the single optimal networks. **D**. Connectivity per site for the MutInf (round 5) selection procedure starting from *F*_*AP C*_, 1*Link, random* initial ranking and the *u*_12_ shared networks. **E**. conservation per site (bits) in the MSA.

**Figure C.18:**
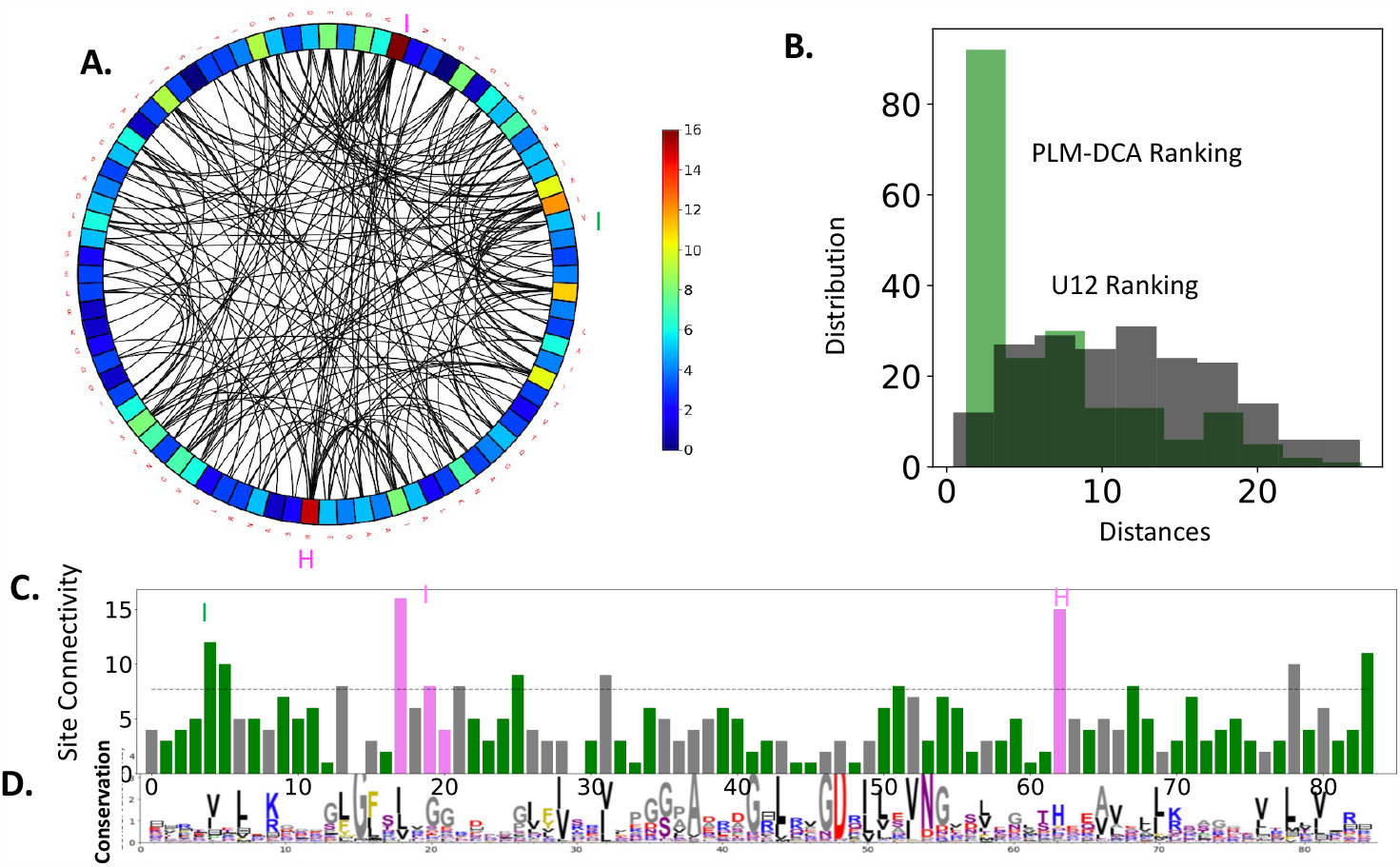
PDZ) Structural and functional Couplings: **A**.: U12 network. The site connectivity is indicated by color-code. **B**. Histogram of distances for the pairs connected in the U12 network, compared with the histogram of distances for the same number of pairs with largest Frobenius Norm with APC correction. Similar histograms are obtained for the single optimal networks. **C**. Connectivity per site for the *u*_12_ shared networks. the most sensitive sites (magenta), and the tolerant sites (green). **D**. Conservation per site (bits) in the MSA.

**Figure C.19:**
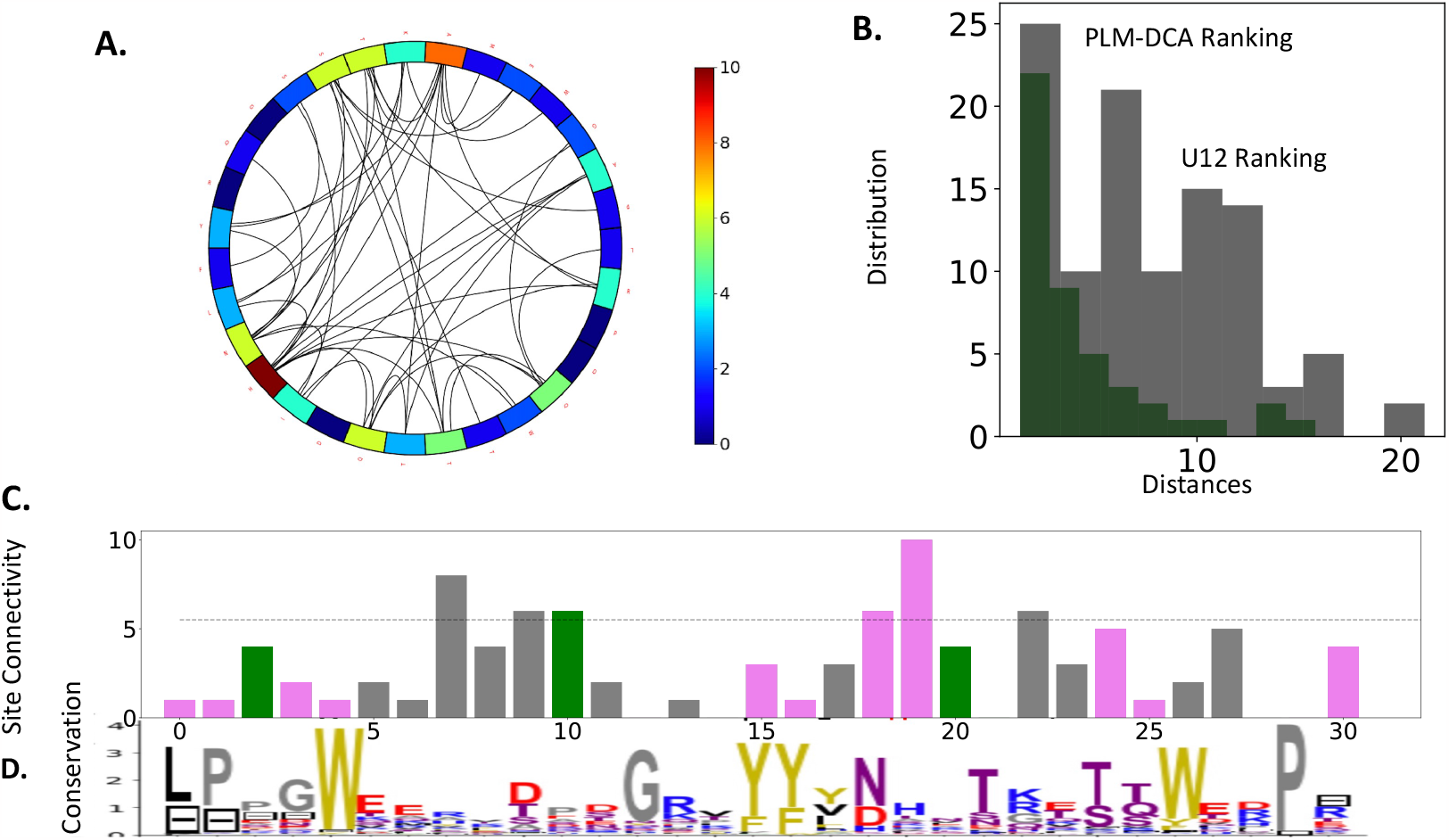
WW) Structural and functional Couplings: **A**.: U12 network. The site connectivity is indicated by color-code. The U12 shared network interconnects sites at or close to the peptide binding surface. The latter is divided in two structurally defined regions: The proline specific binding sites (16Y, 27W) and the specificity sites (18L -20H-25T) in the *β*_2_ strands and the *β*_2_ -*β*_3_ loops. Largely interconnected sites are also in the *β*_1_-*β*_2_ loop agreement with the with the supports [87]) [88]. Moreover, as pointed out in [34] residues 8 and 18, which are also largely interconnected, arbour important stabilizing and functional mutations A8R,L18K. **B**. Histogram of distances for linked pairs in the U12 network, compared with the histogram of distances for the same number of pairs with largest plm DCA Frobenius Norm with APC correction. Similar histograms are obtained for the single optimal networks. Large interconnected sites are not dominated by sites in contacts contrary to what obtained for pairs of sites with large Frobenius norms, but could be interconnected through the peptide. **C**. Connectivity per site for the *u*_12_ shared networks.

## Footnotes

1 The data considered here were obtained at maximal (2,500) ampicillin concentration, see Fig. 6 for comparison with other concentrations.

2 The relative size of the train set is defined with respect to the available experimental single point mutations (Table 1)

3 The contribution to *Sp* due to a new link decreases with the number of links previously added to the network, as the variance of the estimate of the mutational costs increases, see eq:appendix:varh and their dependences on *K*_*i*_.

4 This corresponds to round 0 for heuristic *m*, and round 1 for the other heuristics.

